# Structure-Based Survey of the Human Proteome for Opportunities in Proximity Pharmacology

**DOI:** 10.1101/2022.01.13.475779

**Authors:** Evianne Rovers, Matthieu Schapira

## Abstract

Proximity pharmacology (ProxPharm) is a novel paradigm in drug discovery where a small molecule brings two proteins in close proximity to elicit a signal, generally from one protein onto another. The potential of ProxPharm compounds as a new therapeutic modality is firmly established by proteolysis targeting chimeras (PROTACs) that bring an E3 ubiquitin ligase in proximity to a target protein to induce ubiquitination and subsequent degradation of the target protein. The concept can be expanded to induce other post-translational modifications via the recruitment of different types of protein-modifying enzymes. To survey the human proteome for opportunities in proximity pharmacology, we systematically mapped non-catalytic drug binding pockets on the structure of protein-modifying enzymes available from the Protein Databank. In addition to binding sites exploited by previously reported ProxPharm compounds, we identified putative ligandable non-catalytic pockets in 188 kinases, 42 phosphatases, 26 deubiquitinases, 9 methyltransferases, 7 acetyltransferases, 7 glycosyltransferases, 4 deacetylases, 3 demethylases and 2 glycosidases, including cavities occupied by chemical matter that may serve as starting points for future ProxPharm compounds. This systematic survey confirms that proximity pharmacology is a versatile modality with largely unexplored and promising potential, and reveals novel opportunities to pharmacologically rewire molecular circuitries.

## INTRODUCTION

Proteolysis targeting chimeras (PROTACs) are bifunctional small molecules that simultaneously bind an E3 ubiquitin ligase and a target protein, thereby inducing the ubiquitination and subsequent proteasomal degradation of the protein target^1^. This type of molecules has evolved over the past 20 years from a chemical biology curiosity to a promising therapeutic modality, with clear dosedependent degradation of therapeutic targets such as AR, IRAK4 or BTK observed in man (clinicaltrials.gov identifiers NCT03888612, NCT04772885, NCT04830137), and the question is no longer whether but when the first PROTAC will be approved for therapeutic use by regulatory agencies. With proof-of-concept in sight, the scientific community is now looking at novel ways to apply the concept of proximity pharmacology (ProxPharm), where chemically induced proximity between proteins can be used to rewire the molecular circuitry of cells for chemical biology applications or therapeutic benefit^2,3^. Indeed, ProxPharm compounds were recently reported that recruit a phosphatase, two kinases, an acetyltransferase, and a deubiquitinase to post-translationally modify neo-substrates^4–7^.

Structural studies have shown that PROTACs are not simply acting as chemical linkers but rather stabilize non-natural protein-protein interactions between E3 ligases and target proteins^8^. Because compatible protein interfaces do not always exist between two proteins, a prevailing notion is that a collection of chemical handles binding a diverse array of E3 ligases will be necessary to productively induce the degradation of any given protein. Additionally, the tissue expression profile and subcellular localization of the E3 ligase must match that of the target protein for a PROTAC to be active. Finally, PROTACs recruiting E3 ligases with disease-specific tissue expression profiles can avoid adverse effects associated with the indiscriminate inhibition of the protein target. For example, a senolytic PROTAC exploits the restricted expression profile of the E3 ligase CRBN to avoid toxicity associated with the adverse inhibition of the target protein, Bcl-xl, in platelets^9^. Similar rules are expected to apply to ProxPharm compounds beyond PROTACs, emphasizing the need to identify chemical handles for a diverse array of protein-modifying enzymes.

To uncover novel opportunities for the development of future ProxPharm compounds, we searched for non-catalytic ligandable pockets (structural cavities that can be occupied by small-molecule ligands) in all experimental structures of human protein-modifying enzymes, including kinases, phosphatases, acetyltransferases, deacetylases, methyltransferases, demethylases, glycosyltransferases, glycosidases and deubiquitinases. The ligandability of E3 ligases was previously reviewed^10^ and not considered in this analysis which is focused on opportunities for proximity pharmacology beyond PROTACs^1,10–13^. We identified non-catalytic pockets in 287 human enzymes, including those recruited by previously reported ProxPharm compounds. This analysis further confirms the rich potential of proximity pharmacology for chemical biology applications.

## METHODS

### Mapping binding pockets

A list of enzymes was compiled from the Expasy ENZYME database and the UniprotKB database and mapped to corresponding PDB codes. The 3D structures were extracted from the PDB and the biologically relevant oligomeric state was generated with ICM. The icmPocketfinder module was run against each converted ICM object using default settings. The pockets were categorized as non-catalytic based on the following two approaches.

### Interpro domain analysis

The domain architecture of each enzyme was extracted from the InterPro database^14^. The domains were marked either as catalytic or non-catalytic based on GO ontology or literature. Residues within 2.8Å of the pocket mesh generated by ICM were considered as lining the pocket, and the N- and C-terminal boundaries of this selection were used to define a ‘pseudo’ sequence for the pocket. These sequences were aligned and compared with the domain architecture of the enzyme to determine the domain location of the pocket. If the pocket was in a manually curated non-catalytic domain, the pocket was marked non-catalytic.

### Catalytic residues proximity analysis

For each enzyme, the corresponding catalytic residue information was extracted from either the Mechanism and Catalytic Site Atlas database^15^ or UniprotKB database^16^. If the catalytic residues were present in the structure, the distance between the pockets and the catalytic residues were measured. If the pocket was more than 7 Å away from the catalytic residues, it was categorized as non-catalytic.

### Additional filters

Nucleotide binding residues and co-factor binding residues information was extracted from the UniprotKB database to determine which pockets corresponded to nucleotide or co-factor binding sites. For example, the ATP binding site in protein kinases or the acetyl-CoA binding site in acetyltransferases. If the distance between the pocket and nucleotide/co-factor binding residues was less than 7 Å, the pocket was filtered out. If the pocket was in proximity (<5Å) of unresolved residues in the structure due to poor electron density, the pocket was not included for further analysis. If the catalytic residues were among the missing residues, pockets were excluded as well. Pockets were also excluded when located at the interface of inhibitor proteins and enzyme complexes. Next, pockets were filtered for duplicates (when two structures representing the same enzyme had a similar pocket, the largest pocket was retained) and druggability. Druggability was determined using the pocket properties generated by ICM (volume: 1555.7-661.1 Å^3^, area: 155-655 Å^2^, hydrophobicity: >0.44, buriedness: 0.6-0.95, DLID^17^: >-1). Cut-off values were based on properties of experimentally proven druggable pockets. Lastly, the pockets were grouped based on their domains. A list of manually curated non-catalytic domains was formed, from which non-catalytic domains necessary for the catalytic activity were excluded.

### Cysteine reactivity

The predicted reactivity of cysteine side-chains lining pockets was predicted using the ReactiveCys module of ICM. The method is based on reactivity data for 34 reactive and 184 non-reactive cysteines from isoTOP-ABPP (isotopic tandem orthogonal proteolysis activity-based protein profiling)^18^ and a nonredundant set of PDB protein structures (resolution < 2.5 A) with covalently-modified cysteines (272 reactive).

## RESULTS

To assemble a list of druggable binding pockets that may be exploited by ProxPharm compounds, all high-resolution structures of human protein-modifying enzymes beyond E3 ligases in the PDB were analyzed with the cavity mapping tool IcmPocketFinder (Molsoft, San Diego). Only structural cavities with properties (volume, area, hydrophobicity, buriedness and drug-like density (DLID)) within a pre-defined range (detailed in the Methods section) were deemed ligandable and were considered further. A permissive definition of ligandability was used to reflect the fact that chemical handles for ProxPharm compounds do not need to bind potently to their target. Indeed, ligands with up to 10 μM affinity have been successfully used to make PROTACs^19^. When a ligandable cavity was found in a non-catalytic domain, the domain was also deemed ligandable in the context of enzymes not in the PDB, but with a low confidence score. When enzymes were bound to other proteins in the PDB, cavities were also searched at the protein interface. Pockets that may be exploited by ProxPharm compounds could be divided into three categories: 1) those located in non-catalytic domains, 2) those found at non-catalytic sites of the catalytic domain, 3) those mapping at the interface of protein complexes (Figure 1).

**Figure 1.**
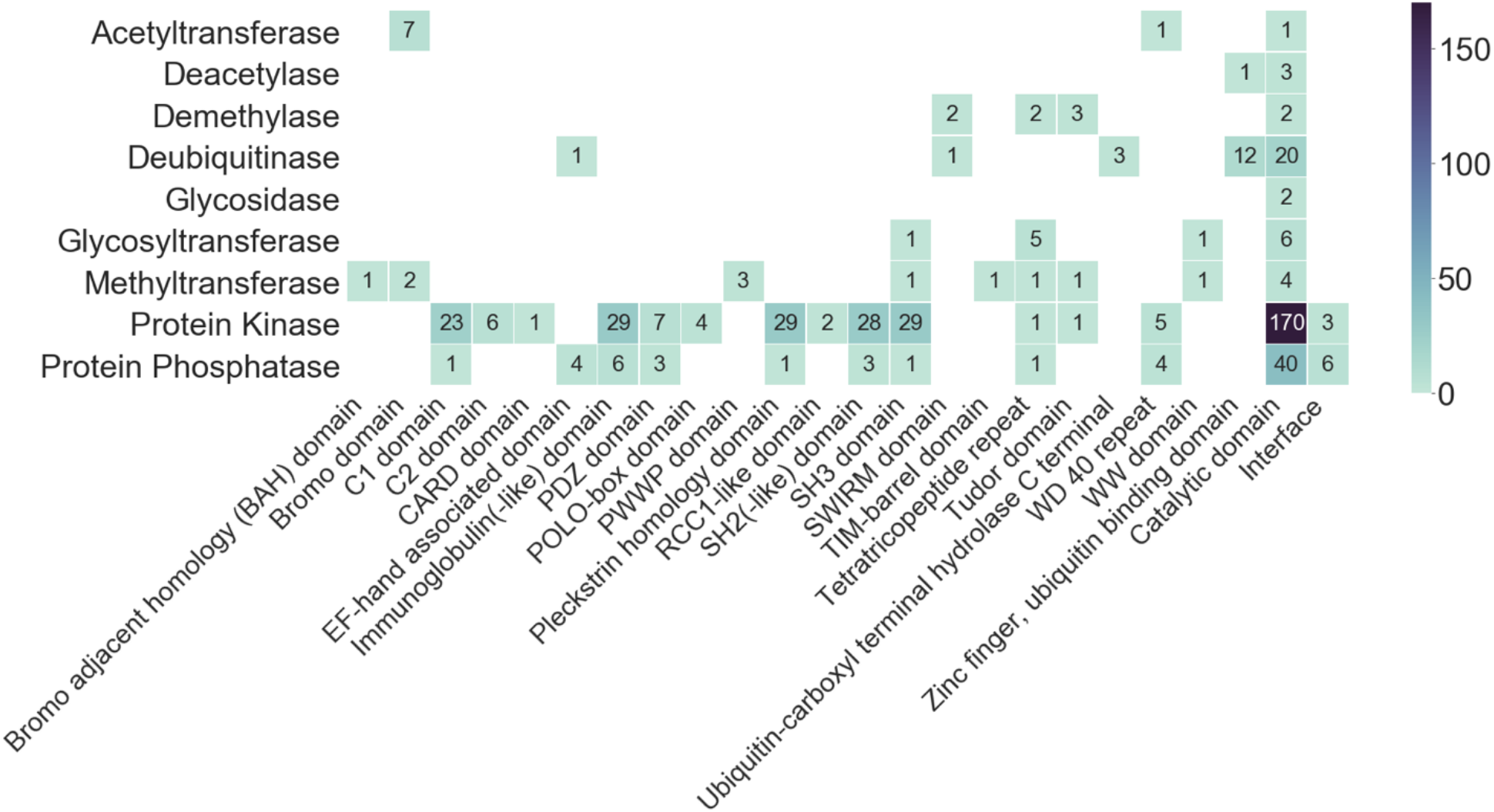
Distribution of non-catalytic pockets in human protein-modifying enzymes. The number of proteins with a putative ligandable non-catalytic pocket is shown for each structural domain and each protein family.

Potentially ligandable non-catalytic pockets were found in 188 kinases, 42 phosphatases, 26 deubiquitinases, as well as several writers and erasers of methyl, acetyl and glycosyl groups (Figure 1, Table S1 and S2). In the following section, we review in detail each protein family.

### Protein kinases

Ligandable non-catalytic pockets were found in the catalytic domain of 170 kinases (Figure 1, Table S2). For instance, in 86 kinases, a pocket is found in the a-lobe of the kinase domain (Figure 2A, Pocket PK3) and, in the context of Abelson kinase, is exploited by an activating compound located over 15Å away from the imatinib-occupied active site (Figure 3, PDB 6NPU)^20^. Other pockets are recurrently found at five other locations and could potentially be exploited to pharmacologically hijack kinases (Figure 2A). In particular, 47 kinases share a cavity below the sub-activation loop (Figure 2A, Pocket PK4) which is occupied by a small molecule in the MAP kinase p38α^21^ (PDB 3HVC). A β-lobe cavity is found in another 25 kinases (Figure 2A, Pocket PK5), where, in PDK1, a cysteine is covalently engaged by fragment inhibitors or activators (PDB 3ORZ)^22^ and a different β-lobe pocket is identified in 9 kinases (Figure 2A, Pocket PK2) and occupied by a fragment molecule in the context of CDK2 (PDB 6Q4D)^23^.

**Figure 2.**
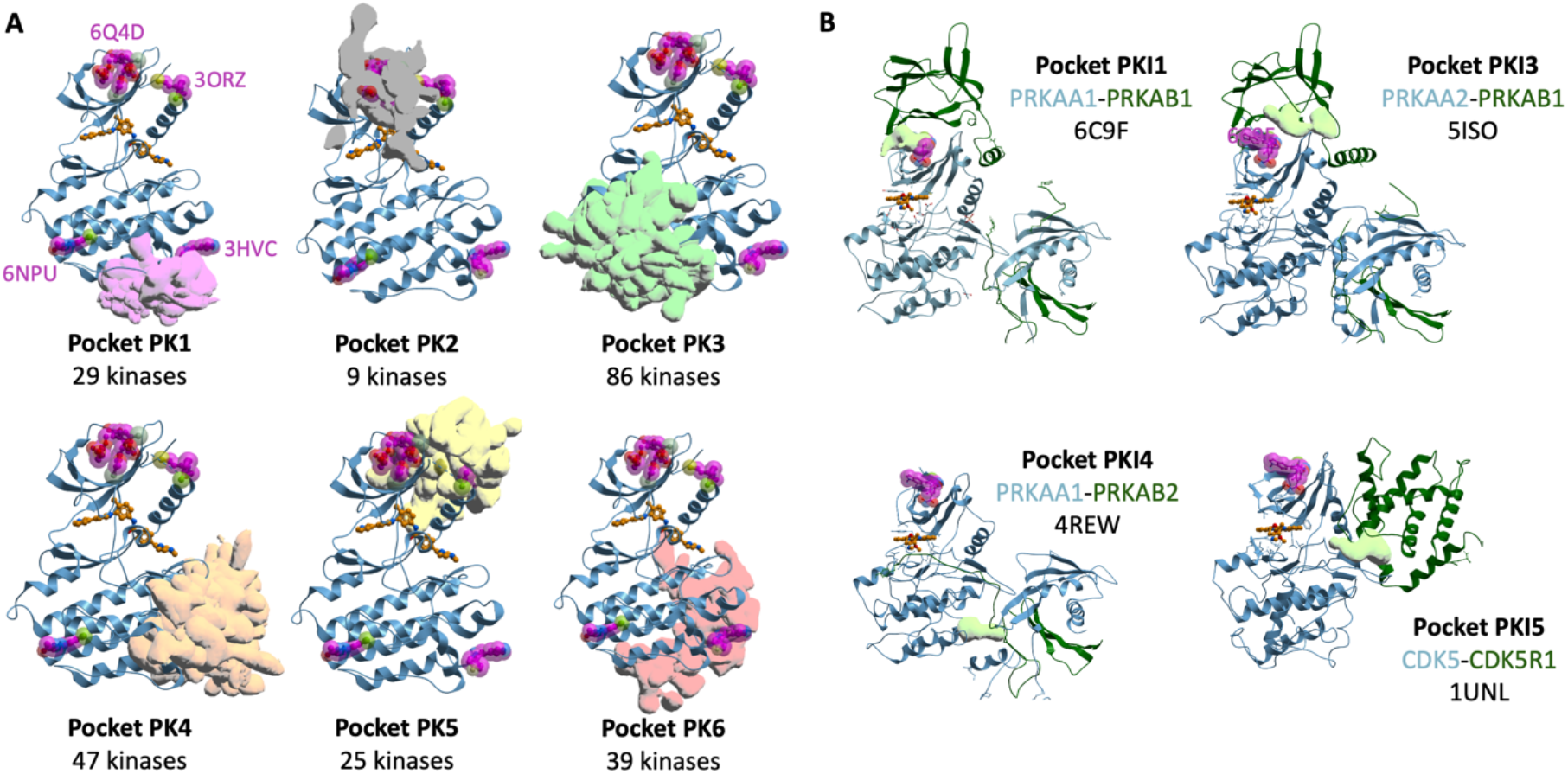
Recurrent non-catalytic pockets mapping at protein kinase domains. A) Pockets found in the kinase domain. ABL1 (blue) bound to catalytic inhibitor (orange) is used as a canonical reference structure (PDB 6NPU^20^). Recurrent pockets are shown as overlapping meshes colored based on their location. Allosteric ligands are shown in purple as references on all structures (PDB 6Q4D^23^, 3ORZ^22^, 6NPU^20^, 3HVC^21^). B) Pockets (lightgreen) found at the interface of the kinase domain (blue) and interacting proteins (darkgreen) in complex structures of PRKAA1 (PDB 6C9F^24^, 4REW^25^), PRKAA2 (PDB 5ISO), and CDK5 (PDB 1UNL^26^). Catalytic and allosteric ligands are shown in orange and purple as a reference on all structures (reference ligands PDB 6C9F^24^).

**Figure 3.**
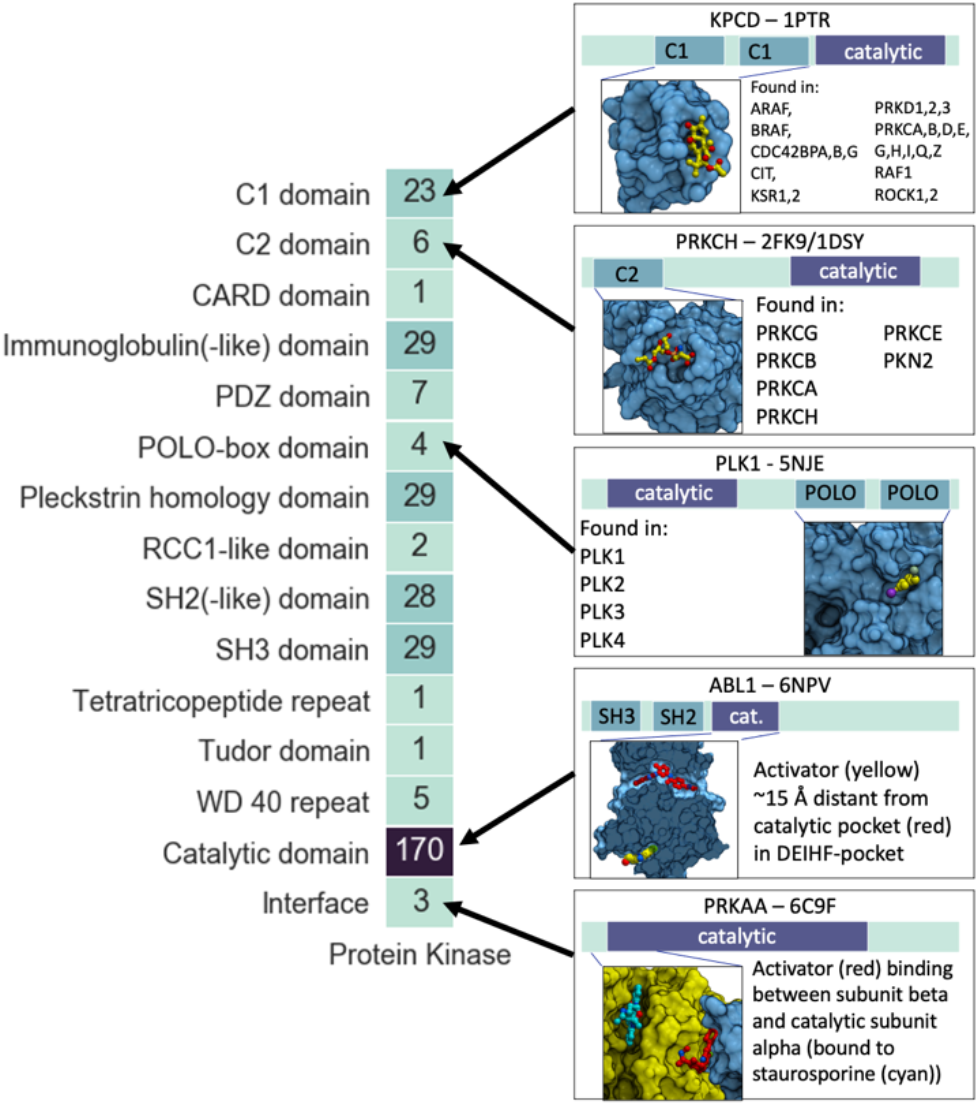
Examples of non-catalytic pockets found in diverse domains of protein kinases. Diacylglycerol bound to the C1 domain of KPCD (PDB 1PTR^33^), phosphatidylserine (PDB 1DSY^34^) bound to the C2 domain of PRKCH (PDB 2FK9^35^), fragment bound to the POLO domain of PLK1 (PDB 5NJE^29^), allosteric activators bound to the catalytic domain of ABL1 (PDB 6NPV^20^) and PRKAA (PDB 6C9F^24^). Details are provided in Table S1.

Ligandable pockets were also found at the interface of the catalytic domain of 3 kinases (PRKAA1, PRKAA2, CDK5) and cofactor proteins (Figure 2B). For example, a pharmacological activator is sandwiched at the interface of the β-lobe of PRKAA1 and its cofactor PRKAB1 (Figure 2B, Pocket PKI1)^24^. Interestingly, this chemical scaffold was recently linked to an inhibitor of Bruton’s tyrosine kinase (BTK), leading to the phosphorylation of BTK by PRKAA1 in cells, in what was the first example of a phosphorylation-inducing chimeric small molecule (PHIC)^4^.

Multiple potentially ligandable cavities were also identified in non-catalytic domains of kinases. For example, a cavity was found in the non-catalytic C1 domain of 23 kinases such as BRAF, CDC42 binding kinases, or PKC kinases (Table S1, Figure 3). Binding of diacylglycerol to this pocket leads to translocation from the cytosol to the membrane of PKC kinases, and catalytic activation^27^. The cavity was successfully targeted by drug-like molecules such as V8-benzolactams^28^, which can be used as PKC-recruiting handles in heterobifunctional PHICS. Using this strategy, Siriwardena et al. could induce the phosphorylation of BRD4 by PKC4^4^.

A membrane-targeting C2 domain is also present in 6 protein kinases, including PKC kinases, but the ligandability of its phosphatidylserine binding pocket is unclear. A tyrosine-lined pocket conserved in the POLO domain of PLK kinases participates in substrate recognition and was targeted by weak compounds that would need to be optimized to serve as ProxPharm handles (Figure 3)^29^. Five kinases contain a WD-40 repeat (WDR), which is a β-propeller domain with a druggable central cavity^30^. For instance, the WDR domain of LRRK2 could be exploited by future PHICS to phosphorylate targets in the brain, where it is expressed.

Other protein domains of potential interest were identified in human kinases, but even though cavities meeting our selection criteria were found, the general ligandability of these domains remains to be supported experimentally. For instance, 29 kinases contain an immunoglobulin-like domain (Figure 1,3). Small molecule ligands were shown to bind to the immunoglobulin-like domain of the unrelated protein RAGE, but ligands were prohibitively weak^31^. Another 28 kinases contain both SH2 and SH3 domains (Figure 1,3), known to participate in the formation of an auto-inhibitory state and contribute to substrate recruitment of Src family kinases. Despite sustained efforts, potent, drug-like, cell-penetrant ligands remain to be found for these domains. Nevertheless, they may be sufficiently ligandable for the discovery of weak compounds that may serve as valid chemical handles for kinase-recruiting ProxPharms. In another example, the poorly characterized kinase STK31 includes a Tudor domain (Table 2, Figure 1), generally found in proteins involved in chromatin-mediated signaling. This domain was targeted by a potent chemical probe in the context of the methyltransferase SETDB1^32^ and may be ligandable in STK31.

### Protein phosphatases

Non-catalytic pockets were found in 43 protein phosphatases (Table 1). Among these, 40 were in the catalytic domain and 24 in juxtaposed domains (Figure 1). Some of the non-catalytic cavities were recurrently found in the phosphatase domain: 14 tyrosine-protein phosphatases share a cavity 15Å from the catalytic site (Figure 4A, Pocket PP3), which, in the context of PTPN5, is occupied by an allosteric activator (PDB 6H8R)^36^. Other recurrent cavities are found at five other locations of the catalytic domain and could potentially be exploited to recruit tyrosine-protein phosphatases to target proteins. Furthermore, 5 serine/threonine-protein phosphatases have 4 recurrent non-catalytic cavities in their catalytic domain (Figure 4B).

**Figure 4.**
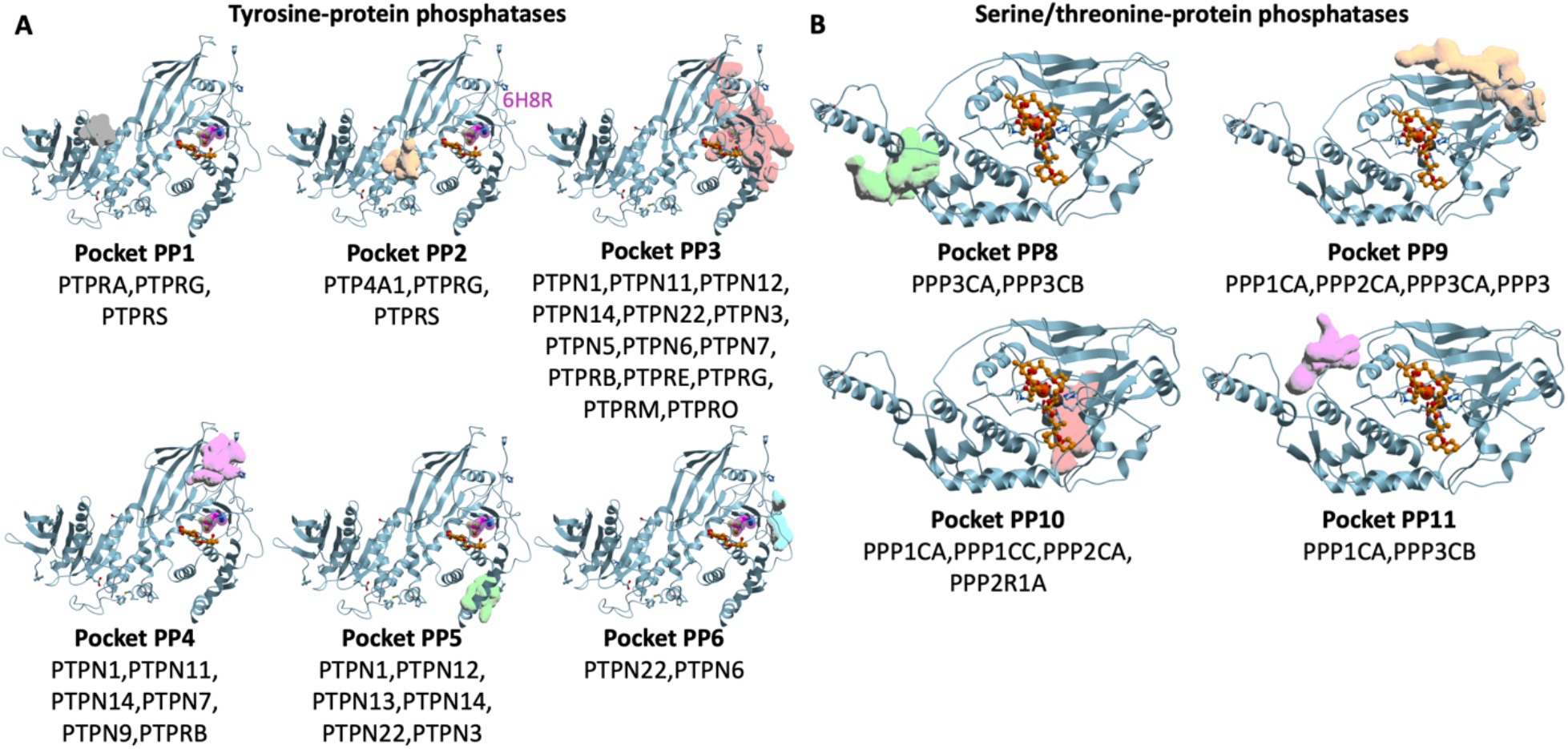
Recurrent non-catalytic pockets in catalytic domain of *protein* phosphatases. A) Tyrosine-protein phosphatases. Reference structure: PDB 2NLK; reference catalytic inhibitor (orange): PDB 1L8G*^37^*, B) Serine/threonine-protein phosphatases. Reference structure: PDB 1AUI*^38^*; reference catalytic inhibitor: PDB 2IE4*^39^*.

Non-catalytic pockets were also found at multiple protein-protein interfaces, including a cavity located at the interface of the three subunits of the protein phosphatase 2A (PP2A) heterotrimer, and occupied by a small molecule activator^40^ (Figure 5, Pocket PPI1). Heterobifunctional compounds derived from this activator could potentially be used for targeted dephosphorylation. This hypothesis is further supported by the fact that PP2A was successfully recruited to dephosphorylate the kinases AKT or EGFR by linking kinase inhibitors to peptidic ligands that exploit the tetratricopeptide repeat domain in PP2A5.

**Figure 5.**
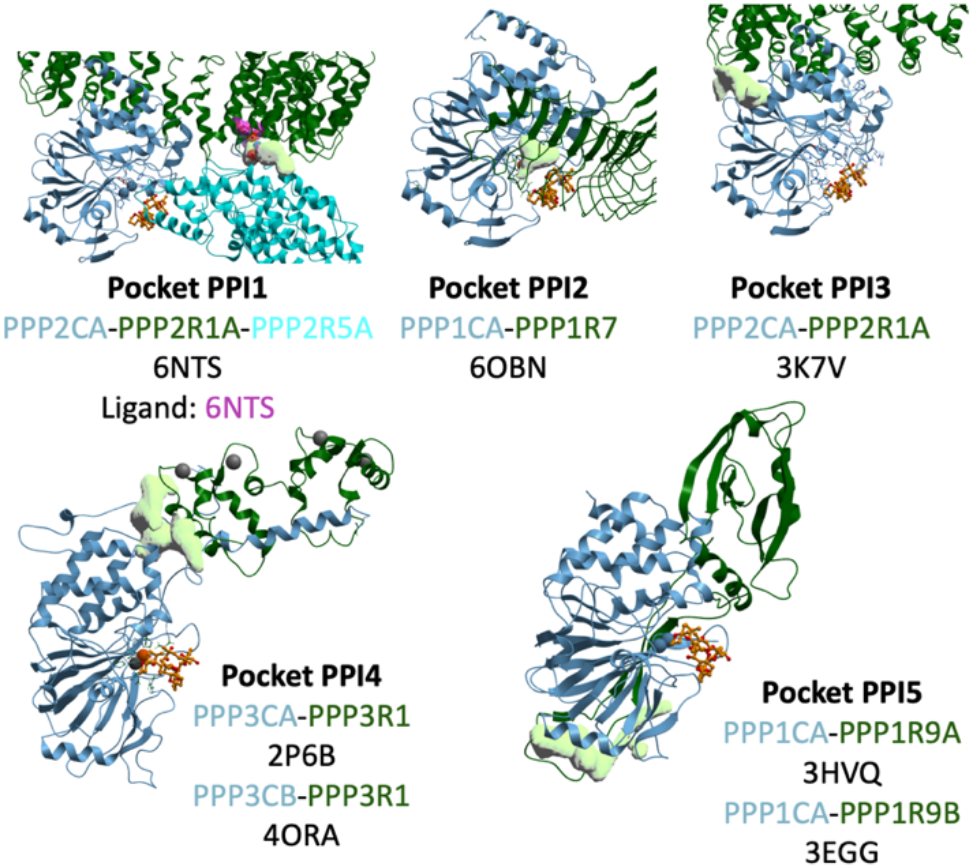
Pockets found at the interface of the protein phosphatase domain (blue) and interacting protein domains (cyan and darkgreen). A catalytic ligand is shown in orange as a reference on all structures (reference ligand PDB 3K7V^41^). Pockets are depicted as green mesh and allosteric activator (PDB 6NTS^40^) is shown in purple.

Cavities are also found in the PDZ domain of protein phosphatases PTPN3, PTPN4 and PTPN14 (Figure 6). The ligandability of these pockets is not experimentally validated, but they are occupied by the C-terminal leucine or valine of pentameric peptide ligands^42,43^, and a similar pocket in the PDZ domain of the unrelated protein PICK1 was crystallized in complex with a small molecule binding with sub-micromolar potency^44^. Finally, pockets with unclear ligandability were found in the SH2 domain of phosphatases PTPN6, PTPN11 and TNS2, and the tetratricopeptide repeat of PPP5C (Figure 6).

**Figure 6.**
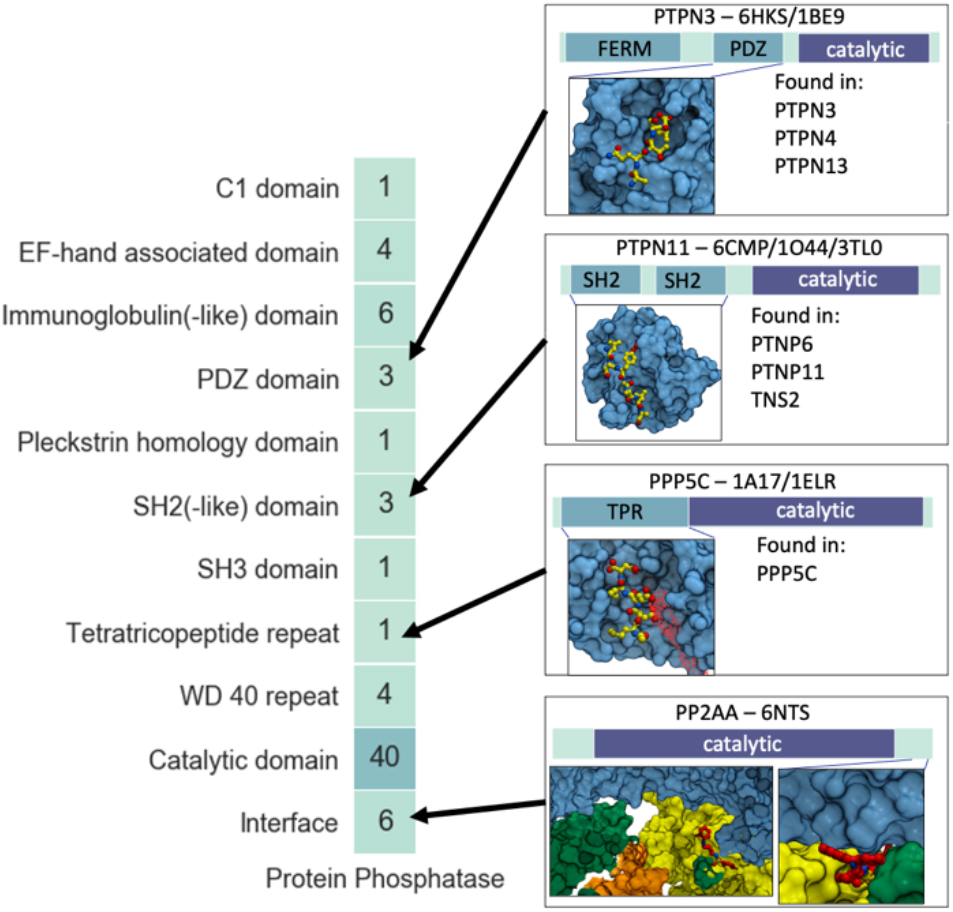
Examples of non-catalytic pockets in protein phosphatases. Pockets in PDZ domain (PDB 6HKS^43^), SH2 domain (PDB 6CMP^45^) and tetratricopeptide repeat (PDB 1A17^46^) with bound peptide (PDB 1BE9^42^, 3TL0^47^ and 1ELR^48^). Allosteric activator for PP2AA (PDB 6NTS^40^) that binds to an allosteric pocket on the interface of the catalytic domain and regulatory proteins. Details are provided in Table S1.

### Protein methyltransferase

Protein methyltransferases (PMTs) are typically large multi-modular proteins where chromatin-binding binding modules are often found juxtaposed to the catalytic domain. For instance, a PWWP domain is found in the NSD subfamily of PMTs (NSD1, NSD2 and NSD3) and chemical probes were reported for the PWWP domain of NSD2 and NSD3 (Figure 7)^56,57^. The NSD3 ligand was recently used as the chemical handle of an NSD3-degrading PROTAC^58^. These ligands – which do not inhibit the enzymatic activity – could also potentially serve as chemical moieties to recruit NSD2 or NSD3 for the methylation of new protein substrates.

**Figure 7.**
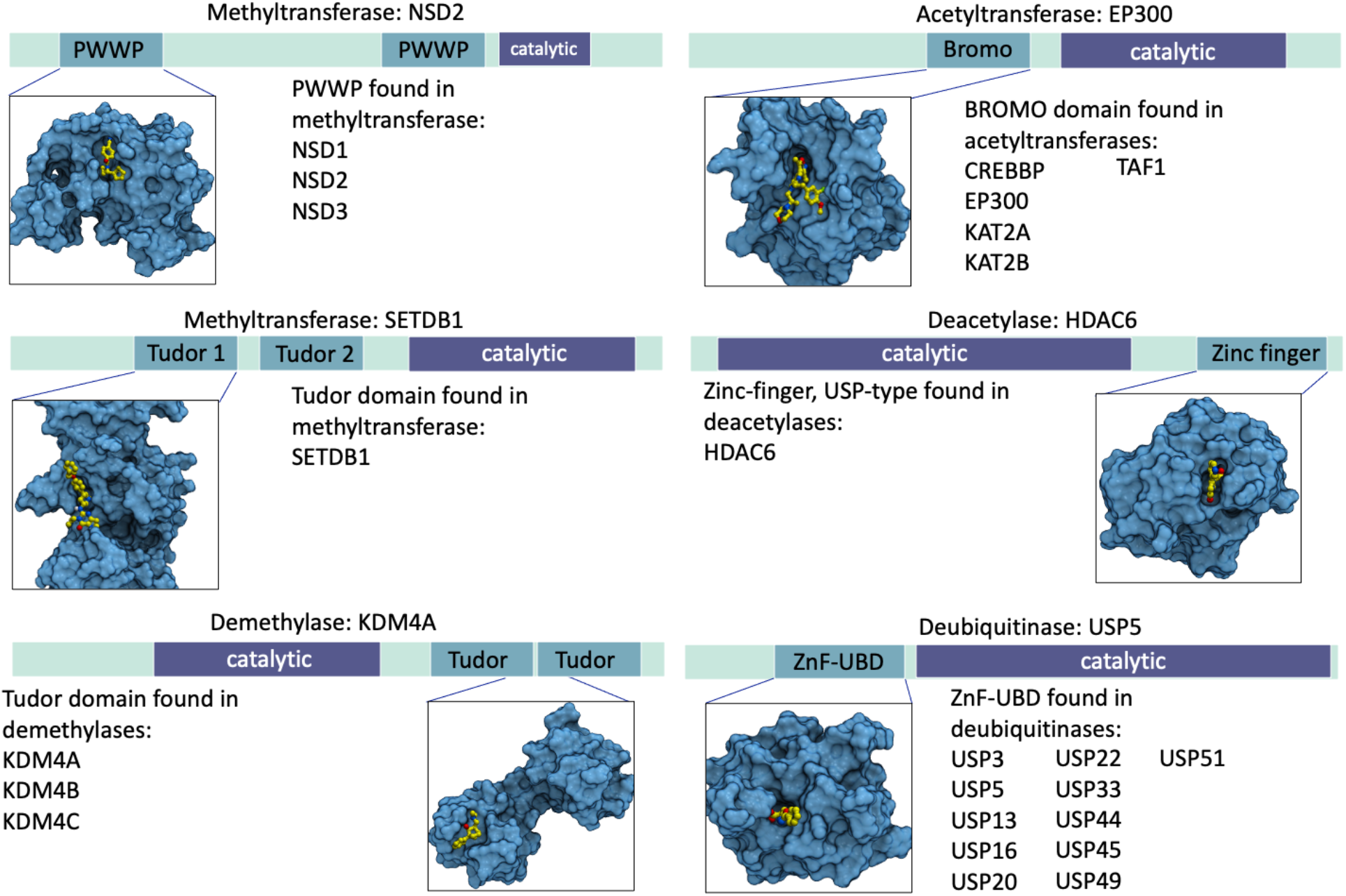
Examples of non-catalytic pockets in non-catalytic domains in methyltransferases, acetyltransferases, demethylases, deacetylases and deubiquitinases. PWWP domain in NSD2 with small-molecule ligand (PDB: 6UE6^49^, pocket PDB: 6XCG), bromo domain in EP300 with small-molecule ligand (PDB: 5BT3, pocket PDB: 6GYT^50^), Tudor domain in SETDB1 with smallmolecule ligand (PDB: 7CJT^32^, pocket PDB: 3DLM), zinc-finger, UBP-type in HDAC6 with small-molecule ligand (PDB: 5KH7^51^, pocket PDB: 5B8D^51^), Tudor domain in KDM4A with chemical fragment (PDB: 5VAR^52^, pocket PDB: 2QQS^53^), ZnF-UBD in USP5 with chemical probe (PDB: 6DXT^54^, pocket PDB: 2G43^55^).

SETDB1, another multi-modular PMT, includes a non-catalytic Tudor domain selectively targeted by a potent chemical probe that may be linked to other ligands to methylate non-natural protein substrates (Figure 7)^32^. Interestingly, recurrent genetic aberrations drive the overexpression of NSD2 in multiple myeloma and pediatric leukemia, and of NSD2, NSD3 and SETDB1 in lung cancer^59–62^, which could possibly offer an opportunity for targeted protein methylation in cells presenting a specific disease-associated genetic profile. Putative ligandable cavities were found in a few other non-catalytic domains of PMTs, including the bromodomain of KMT2A and ASH1L (bromodomains are typically druggable (Figure 1)^63^, but no ligand was reported for these domains.

A recurrent pocket was also found in the catalytic domain of two protein arginine methyltransferases, PRMT3 and PRMT8, which is located more than 17Å away from the catalytic site (Figure S1, Pocket M1). Other unique non-catalytic pockets were found in the methyltransferase domain of 3 PMTs (PRMT3, PRMT5, CARM1) (Table S2). These cavities met our ligandability criteria but so far, their chemical tractability was not validated experimentally.

### Lysine demethylases

A number of non-catalytic domains of lysine demethylases include potentially ligandable pockets. KDM4A, KDM4B and KDM4C all have a Tudor domain, which was shown to be chemically tractable in the context of SETDB1. The Tudor domain of KDM4A was crystallized in complex with a low-affinity chemical fragment (KD~ 80μM) that may be optimized into a stronger-binding chemical handle towards the development of a demethylase-recruiting bifunctional molecules (Figure 7)^52^. Putative ligandable pockets were also found in the tetratricopeptide repeat of KDM6A and UTY and the SWIRM domain of KDM1A and KDM1B (Table S1, Figure 1), but no ligand was so far reported for these domains.

### Lysine acetyltransferases

With over 3000 acetylated lysine side-chains across 1700 human proteins, acetylation is a ubiquitous post-translational modification involved in a diverse array of cellular machineries such as the regulation of gene expression, splicing or cell cycle^64,65^. Out of 35 lysine acetyltransferases in the human genome, we found non-catalytic ligandable pockets in nine (Table S2, Figure 1). Several acetyltransferases include an acetyl-lysine binding bromodomain, five of which were crystallized in complex with multiple small-molecule ligands (EP300, CREBBP, KAT2A, KAT2B and TAF1) (Figure 7)^63^. A compound targeting the bromodomain of one of these, EP300, was chemically linked to an FKBP12-binding molecule to successfully induce the acetylation of FKBP12-fusion proteins by EP300, thereby confirming that acetyltransferases are amenable to proximity pharmacology, and strongly suggesting that bromodomain ligands could be used as chemical handles to recruit other acetyltransferases to neo-substrates^7^.

A WDR domain is also found in GTF3C4, a poorly characterized acetyltransferase (Table S1, Figure 1). The structure of this domain was not experimentally solved, but WDR domains are ligandable in the context of other proteins^30,67^, and this enzyme could potentially be harnessed for targeted acetylation.

### Lysine deacetylases

Deacetylases have a limited number of non-catalytic domains and a ligandable site was found in only one of them: the zinc-finger ubiquitin-binding domain (Znf-UBD) of HDAC6 (Figure 7). This binding pocket recognizes the C-terminal extremity of ubiquitin and was successfully targeted by small molecule ligands^68^ representing excellent chemical handles for proximity pharmacology applications. Non-catalytic pockets were also found in the catalytic domain of three other deacetylases: HDAC4, HDAC8 and HDAC1, but the ligandability of these sites remains to be experimentally validated (Figure 9).

**Figure 9.**
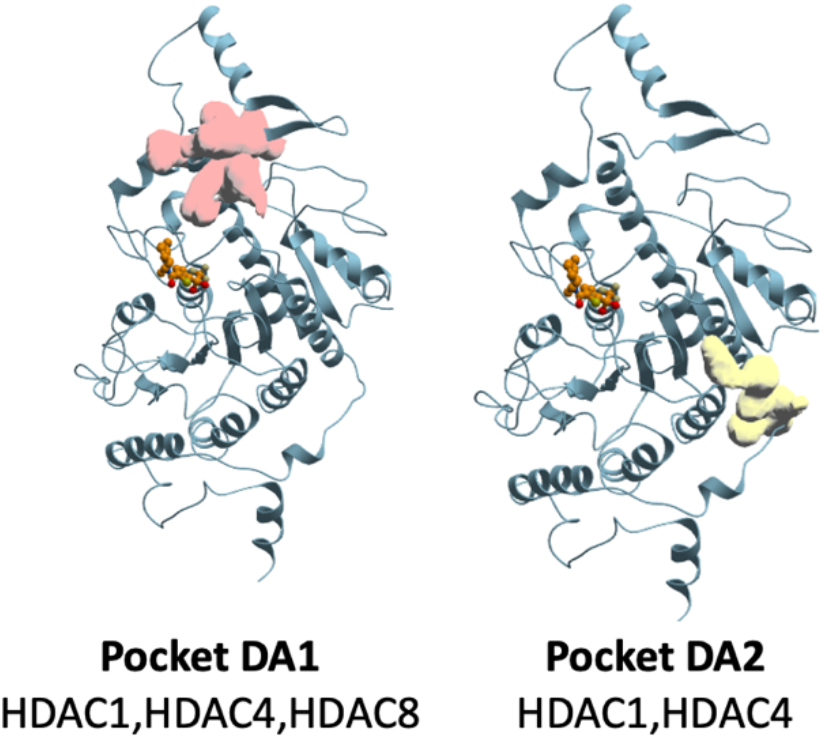
Recurrent non-catalytic pockets in the catalytic domain of histone deacetylases. Lysine deacetylase (blue) bound catalytic inhibitor (orange). Reference structure: PDB 2VQJ*^66^*; reference catalytic inhibitor (orange): PDB 2VQJ*^66^*.

### Deubiquitinases

Deubiquitinases (DUBs) typically remove ubiquitin tags deposited by E3 ligases. When these tags are signalling for proteasomal degradation, DUBs deubiquitinate and rescue their protein substrates from the ubiquitin-proteasome system and have a stabilizing effect on their target. Chemical handles binding non-catalytic pockets of DUBs may therefore enable the recruitment of DUBs for targeted protein stabilization. As a proof-of-concept, a bifunctional molecule linking a ligand that covalently engages the DUB OTUB1 to a chemical moiety that binds ΔF508-CFTR in cystic fibrosis could stabilize ΔF508-CFTR in an OTUB1-dependent manner^6^. There is no structural information on the N-terminal domain of OTUB1 that is covalently recruited by this chimeric compound, but structures of other non-catalytic domains in DUBs reveal other opportunities for targeted protein stabilization.

The most recurrent ligandable non-catalytic domain of DUBs is the Znf-UBD, found in 11 ubiquitin-specific proteases (USPs, a class of DUBs) (Figure 7, Table S1). Low micromolar ligands were reported for the Znf-UBD of USP5, but these compounds were shown to inhibit the catalytic activity of USP5 and therefore cannot be used as chemical handles to productively recruit USP5 to neo-substrates^54,69^. However, the function of the Znf-UBD of DUBs is poorly understood in other USPs, and ligands targeting this domain may still be valid handles for targeted protein stabilization in the context of other DUBs.

Ligandable pockets were also found in a tandem ubiquitin-like domain located at the C-terminus of three DUBs: USP7, 11, 15 (Figure 1, Table S1). In the context of USP7, this domain binds and activates the catalytic domain^73^. In the absence of structure of full-length USP7 in its activated form, it is unclear whether ligands occupying this C-terminal binding pocket would preserve the activation mechanism of USP7 and could be used to productively recruit USP7 for targeted protein stabilization. Other non-catalytic domains present in deubiquitinases are an EF-hand in USP32 and a SWIRM domain in MYSM1. Chemical ligands have not yet been reported for these domains.

Non-catalytic pockets were recurring at six locations of seven USPs within the peptidase C19-type catalytic domain (Figure 10A, Table S1). Another non-catalytic cavity is observed in the peptidase C12-type catalytic domain of UCHL1 and UCHL5 (Figure 10B). As above, the ligandability of these pockets needs to be confirmed experimentally.

**Figure 10.**
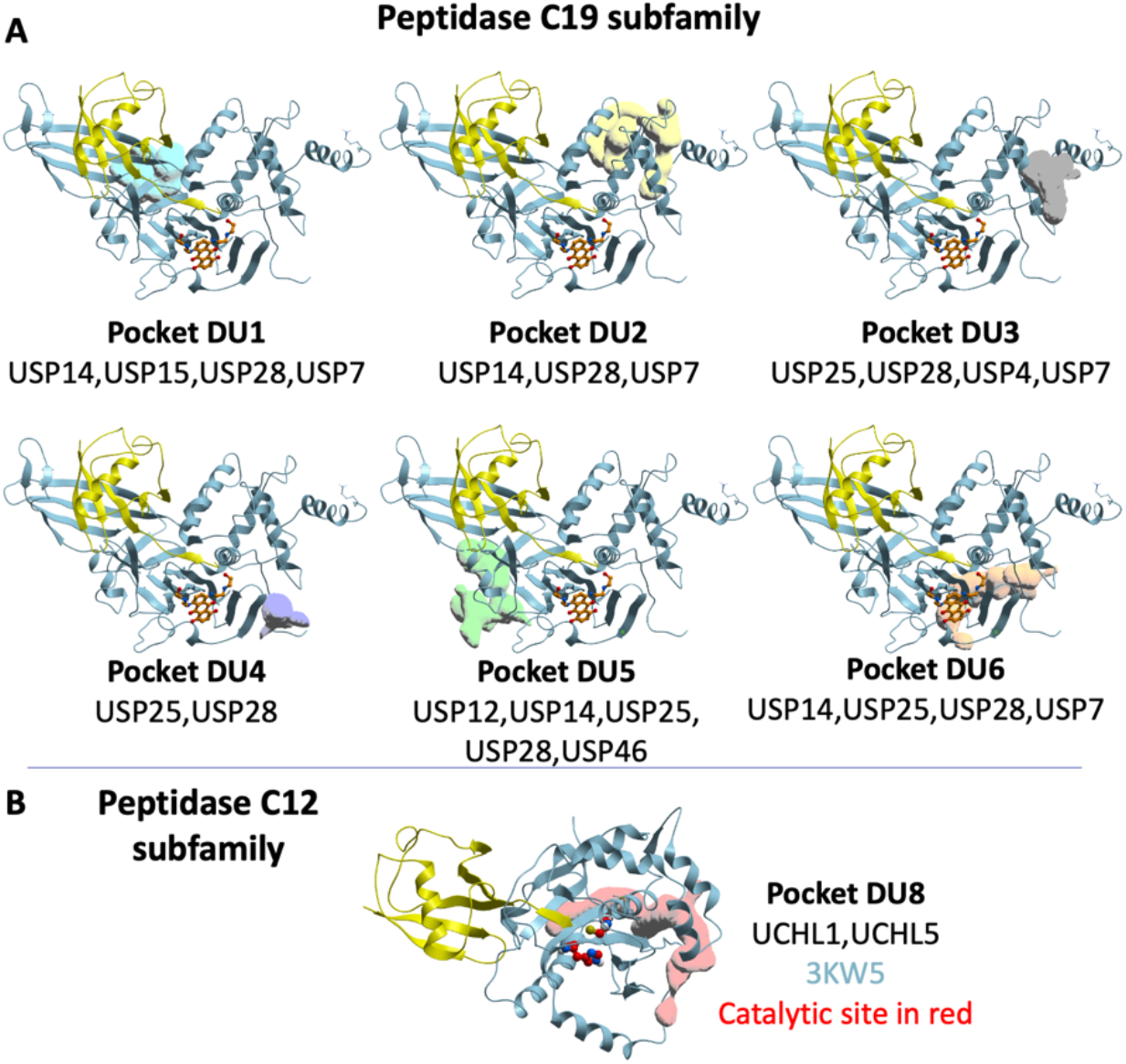
Recurrent non-catalytic pockets in the catalytic domain of peptidases. A) Peptidase C19-type DUBs: reference structure: USP7 (PDB 1BNF^70^), (blue) bound to ubiquitin (yellow). Reference catalytic inhibitor (orange): PDB 6GH9^71^, B) Peptidase C12-type DUBs: UCHL1 (PDB 3KW5^72^) (blue) bound to ubiquitin (yellow). Catalytic residues are highlighted in red.

### Glycosyltransferases

Glycosylation is a post-translational modification that is most common in excreted and extracellular membrane-associated proteins and is frequently dysregulated in diseases, such as cancer or bacterial infection^74^. Proof of principle for proximity-induced glycosylation of target proteins was established by fusing substrate-targeting nanobodies to the glycosyltransferase O-GlcNActransferase (OGT), which effectively induced the glycosylation of the desired protein targets^75^. Putative ligandable pockets in the tetratricopeptide repeat of OGT and TMTC1-4 may be exploited to chemically recruit these glycosyltransferases to neo-substrates. Similarly, the SH3 domain of FUT8 and WW domain of GALNT9 may be considered for the chemical recruitment of these enzymes. Non-catalytic cavities in the glycosyltransferase domain of ST8SIA3, B3GAT1-3, and POFUT2 were also found but, as above, their ligandability should be confirmed experimentally.

### Glycosidases

Similar to glycosyltransferases, protein constructs have been developed using O-GlcNAcase or sialidase connected to nanobody to artificially induce deglycosylation^76–78^. There are limited structures and domain information available for glycosidases, but ligandable pockets are found in the catalytic domain of OGA and MAN1B1 that could be explored for deglycosylation-inducing chimeras.

### Reactive cysteines

PROTACs covalently engaging an E3-ligase have demonstrated that covalent binding is a valid strategy for proximity-induced post-translational modification of target proteins^79–83^. For instance, covalent recruitment of only a small fraction of the cellular pool of the E3-ligase DCAF16 is sufficient to support targeted degradation^80^. A deubiquitinase-targeting chimera also forms a covalent bond with a cysteine of the DUB OTUB16. Electrophylic chemical handles enable the covalent recruitment of domains otherwise not considered ligandable, such as the RING domain of the E3-ligase RNF4^79^, and can be advantageous to enhance potency or selectivity. We used ICM to evaluate the reactivity of cysteine side-chains found in non-catalytic pockets of human protein-modifying enzymes (see Methods section for details).

Reactive cysteines were found in multiple proteins (Table S2). For instance, C576 is lining a pocket in the UBL domain of USP7 C-terminal to the catalytic domain, C210 is found at an ectopic site of the STK16 kinase domain, C266 at a non-catalytic site of the PP2BA phosphatase domain, and C1030 at a cavity remote from the active site of the deacetylase HDAC4 (Figure 11). It would be interesting to screen such proteins with electrophilic fragments to find covalent adducts that may serve as a starting point for novel proximity-pharmacology applications.

**Figure 11.**
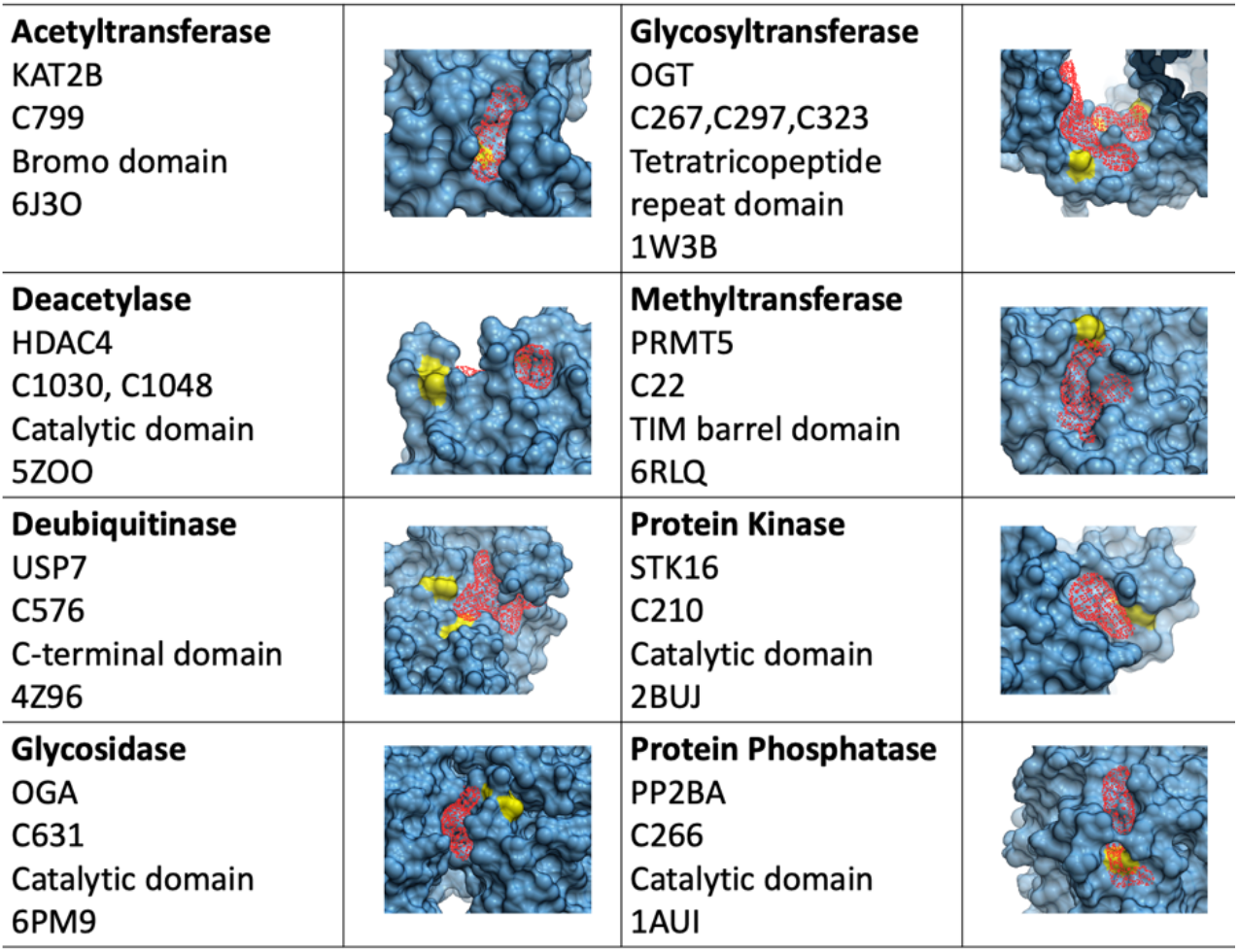
Examples of non-catalytic pockets with reactive cysteine residue lining the cavity. Pockets are highlighted in red. Cysteine residues predicted reactive are colored in yellow.

## DISCUSSION

Our systematic structural survey of the human proteome reveals numerous opportunities for the pharmacological recruitment of protein-modifying enzymes beyond E3 ligases to non-natural substrates. The predicted ligandability of a binding pocket can vary from one method to another and is not a conclusive metric. Here, we use a permissive definition based on volume, area, hydrophobicity, buriedness and DLID values. We first notice that this approach does retrieve binding sites for known ProxPharm compounds, including a protein-protein interface pocket used to recruit the kinase PRKAA (Figure 2B, Pocket PKI1)^4^ and a bromodomain pocket used to recruit the acetyltransferase EP300 (Figure 7)^7^.

Among the collection of binding sites that we compiled, the better validated are the ones for which a high-affinity ligand was already reported (Table S1, confidence level 1). For instance, V8-benzolactams bind the C1 domain of protein kinase C^4,28^, UNC6934 binds the PWWP domain of NSD2^57^ and compound R734 binds a protein interface of the kinase AMPK (Figures 3,7,2)^4,24^. A number of non-catalytic pockets were also found that are targeted by weak ligands that may be valid starting points for the development of ProxPharm compounds (Table S1, confidence level 2). These include compounds and peptides found in the POLO-box domain of PLK1 (Figure 4)^29^ and the PDZ domain of PTPN3 (Figure 6)^42^. Less reliable, but still promising are domains for which no ligand was reported in the context of the protein of interest, but that were shown to be chemically tractable in other proteins (confidence level 3). For example, low nanomolar ligands targeting the WDR domains of EED and WDR5 are in pre-clinical^84–86^ or clinical development (EED clinicaltrials.gov identifier NCT02900651) and WDR domains are found in the kinases LRRK1, LRRK2, MET, MST1R, PIK3R4 and the acetyltransferase GTF3C4 (Table S1). Similarly, Tudor domains are found in demethylases (KDM4A, KDM4B, KDM4C) and protein kinase STK31 (Table S1), and share a canonical aromatic cage with the Tudor domain of SETDB1 targeted by a high-affinity ligand (KD 90 nM)^32^. Finally, sites that meet our ligandability criteria but for which no ligands were found in the protein of interest or close homologues are less reliable, but of potential interest (confidence level 4).

A limitation of our analysis is that we focused exclusively on the structures of enzymes that add or remove chemical or peptidic tags to proteins and are therefore related to E3 ligases in their functional mechanism. In the future, we believe it would be interesting to expand to other enzymes, such as proteases, or potentially to proteins beyond enzymes. Indeed, targeted recruitment of proteins to specific protein interaction hubs may offer novel opportunities to regulate cellular machineries. We also limited our approach to proteins (and homologs) with structural information in the protein databank, but recent breakthroughs in protein structure predictions^87–89^ may enable a future expansion of the analysis to the entire human proteome.

## ASSOCIATED CONTENT

### Supporting Information

The following figures and tables are provided as supporting online information. Putative ligandable domains (defined by their Interpro ID) and associated list of human protein-modifying enzymes (Table S1); Ligandable non-catalytic pockets in protein modifying enzymes including acetyltransferases, deacetylases, methyltransferases, demethylases, glycosyltransferases, glycosidases, deubiquitinases, protein kinases and protein phosphatases (Table S2); Recurrent non-catalytic pockets in catalytic domain of Protein arginine methyltransferases (Figure S1).

## AUTHOR INFORMATION

### Author Contributions

The manuscript was written through contributions of all authors. All authors have given approval to the final version of the manuscript

### Funding Sources

This work was supported by a grant from the Natural Sciences and Engineering Research Council of Canada awarded to MS (grant ALLRP 555329-20). The Structural Genomics Consortium is a registered charity (no: 1097737) that receives funds from Bayer AG, Boehringer Ingelheim, Bristol Myers Squibb, Genentech, Genome Canada through Ontario Genomics Institute [OGI-196], EU/EFPIA/OICR/McGill/KTH/Diamond Innovative Medicines Initiative 2 Joint Undertaking [EUbOPEN grant 875510], Janssen, Merck KGaA (aka EMD in Canada and US), Pfizer and Takeda.

## ABBREVIATIONS

ProxPharm: Proximity Pharmacology
PDB: Protein Data Bank

## Notes

### Competing Interest Statement

The authors have declared no competing interest.

## REFERENCES

(1) Pettersson, M.; Crews, C. M. PROteolysis TArgeting Chimeras (PROTACs) — Past, Present and Future. Drug Discov. Today Technol. 2019, 31, 15–27.

(2) Gerry, C. J.; Schreiber, S. L. Unifying Principles of Bifunctional, Proximity-Inducing Small Molecules. Nat. Chem. Biol. 2020, 16 (4), 369–378.

(3) Conway, S. J. Bifunctional Molecules beyond PROTACs. J. Med. Chem. 2020, 63 (6), 2802–2806.

(4) Siriwardena, S. U.; Munkanatta Godage, D. N. P.; Shoba, V. M.; Lai, S.; Shi, M.; Wu, P.; Chaudhary, S. K.; Schreiber, S. L.; Choudhary, A. Phosphorylation-Inducing Chimeric Small Molecules. J. Am. Chem. Soc. 2020, 142 (33), 14052–14057.

(5) Yamazoe, S.; Tom, J.; Fu, Y.; Wu, W.; Zeng, L.; Sun, C.; Liu, Q.; Lin, J.; Lin, K.; Fairbrother, W. J.; Staben, S. T. Heterobifunctional Molecules Induce Dephosphorylation of Kinases-A Proof of Concept Study. J. Med. Chem. 2020, 63 (6), 2807–2813.

(6) Henning, N. J.; Boike, L.; Spradlin, J. N.; Ward, C. C.; Belcher, B.; Brittain, S. M.; Hesse, M.; Dovala, D.; McGregor, L. M.; McKenna, J. M.; Tallarico, J. A.; Schirle, M.; Nomura, D. K. Deubiquitinase-Targeting Chimeras for Targeted Protein Stabilization. bioRxiv 2021, 2021.04.30.441959.

(7) Wang, W. W.; Chen, L. Y.; Wozniak, J. M.; Jadhav, A. M.; Anderson, H.; Malone, T. E.; Parker, C. G. Targeted Protein Acetylation in Cells Using Heterobifunctional Molecules. J. Am. Chem. Soc. 2021, 143 (40), 16700–16708.

(8) Gadd, M. S.; Testa, A.; Lucas, X.; Chan, K. H.; Chen, W.; Lamont, D. J.; Zengerle, M.; Ciulli, A. Structural Basis of PROTAC Cooperative Recognition for Selective Protein Degradation. Nat. Chem. Biol. 2017 135 2017, 13 (5), 514–521.

(9) He, Y.; Zhang, X.; Chang, J.; Kim, H. N.; Zhang, P.; Wang, Y.; Khan, S.; Liu, X.; Zhang, X.; Lv, D.; Song, L.; Li, W.; Thummuri, D.; Yuan, Y.; Wiegand, J. S.; Ortiz, Y. T.; Budamagunta, V.; Elisseeff, J. H.; Campisi, J.; Almeida, M.; Zheng, G.; Zhou, D. Using Proteolysis-Targeting Chimera Technology to Reduce Navitoclax Platelet Toxicity and Improve Its Senolytic Activity. Nat. Commun. 2020, 11 (1).

(10) Schapira, M.; Calabrese, M. F.; Bullock, A. N.; Crews, C. M. Targeted Protein Degradation: Expanding the Toolbox. Nat. Rev. Drug Discov. 2019, 18 (12), 949–963.

(11) Schneider, M.; Radoux, C. J.; Hercules, A.; Ochoa, D.; Dunham, I.; Zalmas, L. P.; Hessler, G.; Ruf, S.; Shanmugasundaram, V.; Hann, M. M.; Thomas, P. J.; Queisser, M. A.; Benowitz, A. B.; Brown, K.; Leach, A. R. The PROTACtable Genome. Nat. Rev. Drug Discov. 2021, 20 (10), 789–797.

(12) Nalawansha, D. A.; Crews, C. M. PROTACs: An Emerging Therapeutic Modality in Precision Medicine. Cell Chem. Biol. 2020, 27 (8), 998–1014.

(13) Sun, X.; Gao, H.; Yang, Y.; He, M.; Wu, Y.; Song, Y.; Tong, Y.; Rao, Y. PROTACs: Great Opportunities for Academia and Industry. Signal Transduct. Target. Ther. 2019, 4 (1).

(14) Blum, M.; Chang, H. Y.; Chuguransky, S.; Grego, T.; Kandasaamy, S.; Mitchell, A.; Nuka, G.; Paysan-Lafosse, T.; Qureshi, M.; Raj, S.; Richardson, L.; Salazar, G. A.; Williams, L.; Bork, P.; Bridge, A.; Gough, J.; Haft, D. H.; Letunic, I.; Marchler-Bauer, A.; Mi, H.; Natale, D. A.; Necci, M.; Orengo, C. A.; Pandurangan, A. P.; Rivoire, C.; Sigrist, C. J. A.; Sillitoe, I.; Thanki, N.; Thomas, P. D.; Tosatto, S. C. E.; Wu, C. H.; Bateman, A.; Finn, R. D. The InterPro Protein Families and Domains Database: 20 Years On. Nucleic Acids Res. 2021, 49 (D1), D344–D354.

(15) Ribeiro, A. J. M.; Holliday, G. L.; Furnham, N.; Tyzack, J. D.; Ferris, K.; Thornton, J. M. Mechanism and Catalytic Site Atlas (M-CSA): A Database of Enzyme Reaction Mechanisms and Active Sites. Nucleic Acids Res. 2018, 46 (D1), D618–D623.

(16) Bateman, A.; Martin, M. J.; Orchard, S.; Magrane, M.; Agivetova, R.; Ahmad, S.; Alpi, E.; Bowler-Barnett, E. H.; Britto, R.; Bursteinas, B.; Bye-A-Jee, H.; Coetzee, R.; Cukura, A.; Silva, A. Da; Denny, P.; Dogan, T.; Ebenezer, T. G.; Fan, J.; Castro, L. G.; Garmiri, P.; Georghiou, G.; Gonzales, L.; Hatton-Ellis, E.; Hussein, A.; Ignatchenko, A.; Insana, G.; Ishtiaq, R.; Jokinen, P.; Joshi, V.; Jyothi, D.; Lock, A.; Lopez, R.; Luciani, A.; Luo, J.; Lussi, Y.; MacDougall, A.; Madeira, F.; Mahmoudy, M.; Menchi, M.; Mishra, A.; Moulang, K.; Nightingale, A.; Oliveira, C. S.; Pundir, S.; Qi, G.; Raj, S.; Rice, D.; Lopez, M. R.; Saidi, R.; Sampson, J.; Sawford, T.; Speretta, E.; Turner, E.; Tyagi, N.; Vasudev, P.; Volynkin, V.; Warner, K.; Watkins, X.; Zaru, R.; Zellner, H.; Bridge, A.; Poux, S.; Redaschi, N.; Aimo, L.; Argoud-Puy, G.; Auchincloss, A.; Axelsen, K.; Bansal, P.; Baratin, D.; Blatter, M. C.; Bolleman, J.; Boutet, E.; Breuza, L.; Casals-Casas, C.; de Castro, E.; Echioukh, K. C.; Coudert, E.; Cuche, B.; Doche, M.; Dornevil, D.; Estreicher, A.; Famiglietti, M. L.; Feuermann, M.; Gasteiger, E.; Gehant, S.; Gerritsen, V.; Gos, A.; Gruaz-Gumowski, N.; Hinz, U.; Hulo, C.; Hyka-Nouspikel, N.; Jungo, F.; Keller, G.; Kerhornou, A.; Lara, V.; Le Mercier, P.; Lieberherr, D.; Lombardot, T.; Martin, X.; Masson, P.; Morgat, A.; Neto, T. B.; Paesano, S.; Pedruzzi, I.; Pilbout, S.; Pourcel, L.; Pozzato, M.; Pruess, M.; Rivoire, C.; Sigrist, C.; Sonesson, K.; Stutz, A.; Sundaram, S.; Tognolli, M.; Verbregue, L.; Wu, C. H.; Arighi, C. N.; Arminski, L.; Chen, C.; Chen, Y.; Garavelli, J. S.; Huang, H.; Laiho, K.; McGarvey, P.; Natale, D. A.; Ross, K.; Vinayaka, C. R.; Wang, Q.; Wang, Y.; Yeh, L. S.; Zhang, J. UniProt: The Universal Protein Knowledgebase in 2021. Nucleic Acids Res. 2021, 49 (D1), D480–D489.

(17) Sheridan, R. P.; Maiorov, V. N.; Holloway, M. K.; Cornell, W. D.; Gao, Y. D. Drug-like Density: A Method of Quantifying the “Bindability” of a Protein Target Based on a Very Large Set of Pockets and Drug-like Ligands from the Protein Data Bank. J. Chem. Inf. Model. 2010, 50 (11), 2029–2040.

(18) Backus, K. M.; Correia, B. E.; Lum, K. M.; Forli, S.; Horning, B. D.; González-Páez, G. E.; Chatterjee, S.; Lanning, B. R.; Teijaro, J. R.; Olson, A. J.; Wolan, D. W.; Cravatt, B. F. Proteome-Wide Covalent Ligand Discovery in Native Biological Systems. Nature 2016, 534 (7608), 570–574.

(19) Chi, J. (Jack); Li, H.; Zhou, Z.; Izquierdo-Ferrer, J.; Xue, Y.; Wavelet, C. M.; Schiltz, G. E.; Zhang, B.; Cristofanilli, M.; Lu, X.; Bahar, I.; Wan, Y. A Novel Strategy to Block Mitotic Progression for Targeted Therapy. EBioMedicine 2019, 49, 40–54.

(20) Simpson, G. L.; Bertrand, S. M.; Borthwick, J. A.; Campobasso, N.; Chabanet, J.; Chen, S.; Coggins, J.; Cottom, J.; Christensen, S. B.; Dawson, H. C.; Evans, H. L.; Hobbs, A. N.; Hong, X.; Mangatt, B.; Munoz-Muriedas, J.; Oliff, A.; Qin, D.; Scott-Stevens, P.; Ward, P.; Washio, Y.; Yang, J.; Young, R. J. Identification and Optimization of Novel Small C-Abl Kinase Activators Using Fragment and HTS Methodologies. J. Med. Chem. 2019, 62 (4), 2154–2171.

(21) Perry, J. J. P.; Harris, R. M.; Moiani, D.; Olson, A. J.; Tainer, J. A. P38α MAP Kinase C-Terminal Domain Binding Pocket Characterized by Crystallographic and Computational Analyses. J. Mol. Biol. 2009, 391 (1), 1–11.

(22) Sadowsky, J. D.; Burlingame, M. A.; Wolan, D. W.; McClendon, C. L.; Jacobson, M. P.; Wells, J. A. Turning a Protein Kinase on or off from a Single Allosteric Site via Disulfide Trapping. Proc. Natl. Acad. Sci. U. S. A. 2011, 108 (15), 6056–6061.

(23) Wood, D. J.; Lopez-Fernandez, J. D.; Knight, L. E.; Al-Khawaldeh, I.; Gai, C.; Lin, S.; Martin, M. P.; Miller, D. C.; Cano, C.; Endicott, J. A.; Hardcastle, I. R.; Noble, M. E. M.; Waring, M. J. FragLites - Minimal, Halogenated Fragments Displaying Pharmacophore Doublets. An Efficient Approach to Druggability Assessment and Hit Generation. J. Med. Chem. 2019, 62 (7), 3741–3752.

(24) Yan, Y.; Edward Zhou, X.; Novick, S. J.; Shaw, S. J.; Li, Y.; Brunzelle, J. S.; Hitoshi, Y.; Griffin, P. R.; Eric Xu, H.; Melcher, K. Structures of AMP-Activated Protein Kinase Bound to Novel Pharmacological Activators in Phosphorylated, Non-Phosphorylated, and Nucleotide-Free States. J. Biol. Chem. 2019, 294 (3), 953–967.

(25) Li, X.; Wang, L.; Zhou, X. E.; Ke, J.; De Waal, P. W.; Gu, X.; Tan, M. H. E.; Wang, D.; Wu, D.; Xu, H. E.; Melcher, K. Structural Basis of AMPK Regulation by Adenine Nucleotides and Glycogen. Cell Res. 2015 251 2014, 25 (1), 50–66.

(26) Mapelli, M.; Massimiliano, L.; Crovace, C.; Seeliger, M. A.; Tsai, L. H.; Meijer, L.; Musacchio, A. Mechanism of CDK5/P25 Binding by CDK Inhibitors. J. Med. Chem. 2005, 48 (3), 671–679.

(27) Hennings, H.; Blumberg, P. M.; Pettit, G. R.; Herald, C. L.; Shores, R.; Yuspa, S. H. Bryostatin 1, an Activator of Protein Kinase C, Inhibits Tumor Promotion by Phorbol Esters in SENCAR Mouse Skin. Carcinogenesis 1987, 8 (9), 1343–1346.

(28) Mach, U. R.; Lewin, N. E.; Blumberg, P. M.; Kozikowski, A. P. Synthesis and Pharmacological Evaluation of 8-and 9-Substituted Benzolactam-v8 Derivatives as Potent Ligands for Protein Kinase C, a Therapeutic Target for Alzheimer’s Disease. ChemMedChem 2006, 1 (3), 307–314.

(29) Sharma, P.; Mahen, R.; Rossmann, M.; Stokes, J. E.; Hardwick, B.; Huggins, D. J.; Emery, A.; Kunciw, D. L.; Hyvönen, M.; Spring, D. R.; McKenzie, G. J.; Venkitaraman, A. R. A Cryptic Hydrophobic Pocket in the Polo-Box Domain of the Polo-like Kinase PLK1 Regulates Substrate Recognition and Mitotic Chromosome Segregation. Sci. Rep. 2019, 9 (1).

(30) Schapira, M.; Tyers, M.; Torrent, M.; Arrowsmith, C. H. WD40 Repeat Domain Proteins: A Novel Target Class? Nat. Rev. Drug Discov. 2017, 16 (11), 773–786.

(31) Kozlyuk, N.; Gilston, B. A.; Salay, L. E.; Gogliotti, R. D.; Christov, P. P.; Kim, K.; Ovee, M.; Waterson, A. G.; Chazin, W. J. A Fragment-Based Approach to Discovery of Receptor for Advanced Glycation End Products Inhibitors. Proteins 2021, 89 (11), 1399–1412.

(32) Guo, Y.; Mao, X.; Xiong, L.; Xia, A.; You, J.; Lin, G.; Wu, C.; Huang, L.; Wang, Y.; Yang, S. Structure-Guided Discovery of a Potent and Selective Cell-Active Inhibitor of SETDB1 Tudor Domain. Angew. Chem. Int. Ed. Engl. 2021, 60 (16), 8760–8765.

(33) Zhang, G.; Kazanietz, M. G.; Blumberg, P. M.; Hurley, J. H. Crystal Structure of the Cys2 Activator-Binding Domain of Protein Kinase C8 in Complex with Phorbol Ester. Cell 1995, 81, 917–924.

(34) Verdaguer, N.; Corbalan-Garcia, S.; Ochoa, W. F.; Fita, I.; Gómez-Fernández, J. C. Ca2+ Bridges the C2 Membrane-Binding Domain of Protein Kinase Cα Directly to Phosphatidylserine. EMBO J. 1999, 18 (22), 6329–6338.

(35) Littler, D. R.; Walker, J. R.; She, Y. M.; Finerty, P. J.; Newman, E. M.; Dhe-Paganon, S. Structure of Human Protein Kinase C Eta (PKCη) C2 Domain and Identification of Phosphorylation Sites. Biochem. Biophys. Res. Commun. 2006, 349 (4), 1182–1189.

(36) Tautermann, C. S.; Binder, F.; Büttner, F. H.; Eickmeier, C.; Fiegen, D.; Gross, U.; Grundl, M. A.; Heilker, R.; Hobson, S.; Hoerer, S.; Luippold, A.; Mack, V.; Montel, F.; Peters, S.; Bhattacharya, S.; Vaidehi, N.; Schnapp, G.; Thamm, S.; Zeeb, M. Allosteric Activation of Striatal-Enriched Protein Tyrosine Phosphatase (STEP, PTPN5) by a Fragment-like Molecule. J. Med. Chem. 2019, 62 (1), 306–316.

(37) Iversen, L. F.; Andersen, H. S.; Møller, K. B.; Olsen, O. H.; Peters, G. H.; Branner, S.; Mortensen, S. B.; Hansen, T. K.; Lau, J.; Ge, Y.; Holsworth, D. D.; Newman, M. J.; Møller, N. P. H. Steric Hindrance as a Basis for Structure-Based Design of Selective Inhibitors of Protein-Tyrosine Phosphatases†. Biochemistry 2001, 40 (49), 14812–14820.

(38) Kissinger, C. R.; Parge, H. E.; Knighton, D. R.; Lewis, C. T.; Pelletier, L. A.; Tempczyk, A.; Kalish, V. J.; Tucker, K. D.; Showalter, R. E.; Moomaw, E. W.; Gastinel, L. N.; Habuka, N.; Chen, X.; Maldonado, F.; Barker, J. E.; Bacquet, R.; Villafranca, J. E. Crystal Structures of Human Calcineurin and the Human FKBP12–FK506–Calcineurin Complex. Nat. 1995 3786557 1995, 378 (6557), 641–644.

(39) Xing, Y.; Xu, Y.; Chen, Y.; Jeffrey, P. D.; Chao, Y.; Lin, Z.; Li, Z.; Strack, S.; Stock, J. B.; Shi, Y. Structure of Protein Phosphatase 2A Core Enzyme Bound to Tumor-Inducing Toxins. Cell 2006, 127 (2), 341–353.

(40) Leonard, D.; Huang, W.; Izadmehr, S.; O’Connor, C. M.; Wiredja, D. D.; Wang, Z.; Zaware, N.; Chen, Y.; Schlatzer, D. M.; Kiselar, J.; Vasireddi, N.; Schüchner, S.; Perl, A. L.; Galsky, M. D.; Xu, W.; Brautigan, D. L.; Ogris, E.; Taylor, D. J.; Narla, G. Selective PP2A Enhancement through Biased Heterotrimer Stabilization. Cell 2020, 181 (3), 688–701.e16.

(41) Huhn, J.; Jeffrey, P. D.; Larsen, K.; Rundberget, T.; Rise, F.; Cox, N. R.; Arcus, V.; Shi, Y.; Miles, C. O. A Structural Basis for the Reduced Toxicity of Dinophysistoxin-2. Chem. Res. Toxicol. 2009, 22 (11), 1782–1786.

(42) Doyle, D. A.; Lee, A.; Lewis, J.; Kim, E.; Sheng, M.; MacKinnon, R. Crystal Structures of a Complexed and Peptide-Free Membrane Protein-Binding Domain: Molecular Basis of Peptide Recognition by PDZ. Cell 1996, 85 (7), 1067–1076.

(43) Genera, M.; Samson, D.; Raynal, B.; Haouz, A.; Baron, B.; Simenel, C.; Guerois, R.; Wolff, N.; Caillet-Saguy, C. Structural and Functional Characterization of the PDZ Domain of the Human Phosphatase PTPN3 and Its Interaction with the Human Papillomavirus E6 Oncoprotein. Sci. Rep. 2019, 9 (1).

(44) Lin, E. Y. S.; Silvian, L. F.; Marcotte, D. J.; Banos, C. C.; Jow, F.; Chan, T. R.; Arduini, R. M.; Qian, F.; Baker, D. P.; Bergeron, C.; Hession, C. A.; Huganir, R. L.; Borenstein, C. F.; Enyedy, I.; Zou, J.; Rohde, E.; Wittmann, M.; Kumaravel, G.; Rhodes, K. J.; Scannevin, R. H.; Dunah, A. W.; Guckian, K. M. Potent PDZ-Domain PICK1 Inhibitors That Modulate Amyloid Beta-Mediated Synaptic Dysfunction. Sci. Rep. 2018, 8 (1).

(45) Pádua, R. A. P.; Sun, Y.; Marko, I.; Pitsawong, W.; Stiller, J. B.; Otten, R.; Kern, D. Mechanism of Activating Mutations and Allosteric Drug Inhibition of the Phosphatase SHP2. Nat. Commun. 2018 91 2018, 9 (1), 1–14.

(46) Das, A. K.; Cohen, P. T. W.; Barford, D. The Structure of the Tetratricopeptide Repeats of Protein Phosphatase 5: Implications for TPR-Mediated Protein–Protein Interactions. EMBO J. 1998, 17 (5), 1192–1199.

(47) Zhang, Y.; Zhang, J.; Yuan, C.; Hard, R. L.; Park, I. H.; Li, C.; Bell, C.; Pei, D. Simultaneous Binding of Two Peptidyl Ligands by a Src Homology 2 Domain. Biochemistry 2011, 50 (35), 7637–7646.

(48) Scheufler, C.; Brinker, A.; Bourenkov, G.; Pegoraro, S.; Moroder, L.; Bartunik, H.; Hartl, F. U.; Moarefi, I. Structure of TPR Domain–Peptide Complexes: Critical Elements in the Assembly of the Hsp70–Hsp90 Multichaperone Machine. Cell 2000, 101 (2), 199–210.

(49) Ferreira De Freitas, R.; Liu, Y.; Szewczyk, M. M.; Mehta, N.; Li, F.; McLeod, D.; Zepeda-Velázquez, C.; Dilworth, D.; Hanley, R. P.; Gibson, E.; Brown, P. J.; Al-Awar, R.; James, L. I.; Arrowsmith, C. H.; Barsyte-Lovejoy, D.; Min, J.; Vedadi, M.; Schapira, M.; Allali-Hassani, A. Discovery of Small-Molecule Antagonists of the PWWP Domain of NSD2. J. Med. Chem. 2021, 64 (3), 1584–1592.

(50) Ortega, E.; Rengachari, S.; Ibrahim, Z.; Hoghoughi, N.; Gaucher, J.; Holehouse, A. S.; Khochbin, S.; Panne, D. Transcription Factor Dimerization Activates the P300 Acetyltransferase. Nat. 2018 5627728 2018, 562 (7728), 538–544.

(51) Harding, R. J.; Ferreira De Freitas, R.; Collins, P.; Franzoni, I.; Ravichandran, M.; Ouyang, H.; Juarez-Ornelas, K. A.; Lautens, M.; Schapira, M.; Von Delft, F.; Santhakumar, V.; Arrowsmith, C. H. Small Molecule Antagonists of the Interaction between the Histone Deacetylase 6 Zinc-Finger Domain and Ubiquitin. J. Med. Chem. 2017, 60 (21), 9090–9096.

(52) Upadhyay, A. K.; Judge, R. A.; Li, L.; Pithawalla, R.; Simanis, J.; Bodelle, P. M.; Marin, V. L.; Henry, R. F.; Petros, A. M.; Sun, C. Targeting Lysine Specific Demethylase 4A (KDM4A) Tandem TUDOR Domain - A Fragment Based Approach. Bioorg. Med. Chem. Lett. 2018, 28 (10), 1708–1713.

(53) Lee, J.; Thompson, J. R.; Botuyan, M. V.; Mer, G. Distinct Binding Modes Specify the Recognition of Methylated Histones H3K4 and H4K20 by JMJD2A-Tudor. Nat. Struct. Mol. Biol. 2008, 15 (1), 109–111.

(54) Mann, M. K.; Franzoni, I.; De Freitas, R. F.; Tempel, W.; Houliston, S.; Smith, L.; Vedadi, M.; Arrowsmith, C. H.; Harding, R. J.; Schapira, M. Discovery of Small Molecule Antagonists of the USP5 Zinc Finger Ubiquitin-Binding Domain. J. Med. Chem. 2019, 62 (22), 10144–10155.

(55) Reyes-Turcu, F. E.; Horton, J. R.; Mullally, J. E.; Heroux, A.; Cheng, X.; Wilkinson, K. D. The Ubiquitin Binding Domain ZnF UBP Recognizes the C-Terminal Diglycine Motif of Unanchored Ubiquitin. Cell 2006, 124 (6), 1197–1208.

(56) Böttcher, J.; Dilworth, D.; Reiser, U.; Neumüller, R. A.; Schleicher, M.; Petronczki, M.; Zeeb, M.; Mischerikow, N.; Allali-Hassani, A.; Szewczyk, M. M.; Li, F.; Kennedy, S.; Vedadi, M.; Barsyte-Lovejoy, D.; Brown, P. J.; Huber, K. V. M.; Rogers, C. M.; Wells, C. I.; Fedorov, O.; Rumpel, K.; Zoephel, A.; Mayer, M.; Wunberg, T.; Böse, D.; Zahn, S.; Arnhof, H.; Berger, H.; Reiser, C.; Hörmann, A.; Krammer, T.; Corcokovic, M.; Sharps, B.; Winkler, S.; Häring, D.; Cockcroft, X. L.; Fuchs, J. E.; Müllauer, B.; Weiss-Puxbaum, A.; Gerstberger, T.; Boehmelt, G.; Vakoc, C. R.; Arrowsmith, C. H.; Pearson, M.; McConnell, D. B. Fragment-Based Discovery of a Chemical Probe for the PWWP1 Domain of NSD3. Nat. Chem. Biol. 2019, 15 (8), 822–829.

(57) Dilworth, D.; Hanley, R. P.; Ferreira de Freitas, R.; Allali-Hassani, A.; Zhou, M.; Mehta, N.; Marunde, M. R.; Ackloo, S.; Carvalho Machado, R. A.; Khalili Yazdi, A.; Owens, D. D. G.; Vu, V.; Nie, D. Y.; Alqazzaz, M.; Marcon, E.; Li, F.; Chau, I.; Bolotokova, A.; Qin, S.; Lei, M.; Liu, Y.; Szewczyk, M. M.; Dong, A.; Kazemzadeh, S.; Abramyan, T.; Popova, I. K.; Hall, N. W.; Meiners, M. J.; Cheek, M. A.; Gibson, E.; Kireev, D.; Greenblatt, J. F.; Keogh, M.-C.; Min, J.; Brown, P. J.; Vedadi, M.; Arrowsmith, C. H.; Barsyte-Lovejoy, D.; James, L. I.; Schapira, M. A Chemical Probe Targeting the PWWP Domain Alters NSD2 Nucleolar Localization. Nat. Chem. Biol. 2021.

(58) Xu, C.; Meng, F.; Park, K.-S.; Storey, A. J.; Gong, W.; Tsai, Y.-H.; Gibson, E.; Byrum, S. D.; Li, D.; Edmondson, R. D.; Mackintosh, S. G.; Vedadi, M.; Cai, L.; Tackett, A. J.; Kaniskan, H. Ü.; Jin, J.; Wang, G. G. A NSD3-Targeted PROTAC Suppresses NSD3 and CMyc Oncogenic Nodes in Cancer Cells. Cell Chem. Biol. 2021.

(59) Keats, J. J.; Reiman, T.; Maxwell, C. A.; Taylor, B. J.; Larratt, L. M.; Mant, M. J.; Belch, A. R.; Pilarski, L. M. In Multiple Myeloma, t(4;14)(P16;Q32) Is an Adverse Prognostic Factor Irrespective of FGFR3 Expression. Blood 2003, 101 (4), 1520–1529.

(60) Yuan, G.; Flores, N. M.; Hausmann, S.; Lofgren, S. M.; Kharchenko, V.; Angulo-Ibanez, M.; Sengupta, D.; Lu, X.; Czaban, I.; Azhibek, D.; Vicent, S.; Fischle, W.; Jaremko, M.; Fang, B.; Wistuba, I. I.; Chua, K. F.; Roth, J. A.; Minna, J. D.; Shao, N. Y.; Jaremko, Ł.; Mazur, P. K.; Gozani, O. Elevated NSD3 Histone Methylation Activity Drives Squamous Cell Lung Cancer. Nature 2021, 590 (7846), 504–508.

(61) Rodriguez-Paredes, M.; Martinez De Paz, A.; Simó-Riudalbas, L.; Sayols, S.; Moutinho, C.; Moran, S.; Villanueva, A.; Vázquez-Cedeira, M.; Lazo, P. A.; Carneiro, F.; Moura, C. S.; Vieira, J.; Teixeira, M. R.; Esteller, M. Gene Amplification of the Histone Methyltransferase SETDB1 Contributes to Human Lung Tumorigenesis. Oncogene 2014, 33 (21), 2807–2813.

(62) Sengupta, D.; Zeng, L.; Li, Y.; Hausmann, S.; Ghosh, D.; Yuan, G.; Nguyen, T. N.; Lyu, R.; Caporicci, M.; Morales Benitez, A.; Coles, G. L.; Kharchenko, V.; Czaban, I.; Azhibek, D.; Fischle, W.; Jaremko, M.; Wistuba, I. I.; Sage, J.; Jaremko, Ł.; Li, W.; Mazur, P. K.; Gozani, O. NSD2 Dimethylation at H3K36 Promotes Lung Adenocarcinoma Pathogenesis. Mol. Cell 2021, 81 (21).

(63) Filippakopoulos, P.; Knapp, S. Targeting Bromodomains: Epigenetic Readers of Lysine Acetylation. Nat. Rev. Drug Discov. 2014, 13 (5), 337–356.

(64) Verdin, E.; Ott, M. 50 Years of Protein Acetylation: From Gene Regulation to Epigenetics, Metabolism and Beyond. Nat. Rev. Mol. Cell Biol. 2015, 16 (4), 258–264.

(65) Choudhary, C.; Kumar, C.; Gnad, F.; Nielsen, M. L.; Rehman, M.; Walther, T. C.; Olsen, J. V.; Mann, M. Lysine Acetylation Targets Protein Complexes and Co-Regulates Major Cellular Functions. Science 2009, 325 (5942), 834–840.

(66) Bottomley, M. J.; Lo Surdo, P. Lo; Di Giovine, P. Di; Cirillo, A.; Scarpelli, R.; Ferrigno, F.; Jones, P.; Neddermann, P.; De Francesco, R.; Steinkühler, C.; Gallinari, P.; Carfí, A. Structural and Functional Analysis of the Human HDAC4 Catalytic Domain Reveals a Regulatory Structural Zinc-Binding Domain. J. Biol. Chem. 2008, 283 (39), 26694–26704.

(67) Song, R.; Wang, Z. D.; Schapira, M. Disease Association and Druggability of WD40 Repeat Proteins. J. Proteome Res. 2017, 16 (10), 3766–3773.

(68) Ferreira De Freitas, R.; Harding, R. J.; Franzoni, I.; Ravichandran, M.; Mann, M. K.; Ouyang, H.; Lautens, M.; Santhakumar, V.; Arrowsmith, C. H.; Schapira, M. Identification and Structure-Activity Relationship of HDAC6 Zinc-Finger Ubiquitin Binding Domain Inhibitors. J. Med. Chem. 2018, 61 (10), 4517–4527.

(69) Mann, M. K.; Zepeda-Velázquez, C. A.; González-Álvarez, H.; Dong, A.; Kiyota, T.; Aman, A. M.; Loppnau, P.; Li, Y.; Wilson, B.; Arrowsmith, C. H.; Al-Awar, R.; Harding, R. J.; Schapira, M. Structure-Activity Relationship of USP5 Inhibitors. J. Med. Chem. 2021, 64 (20), 15017–15036.

(70) Clarke, J.; Henrick, K.; Fersht, A. R. Disulfide Mutants of Barnase I: Changes in Stability and Structure Assessed by Biophysical Methods and X-Ray Crystallography. J. Mol. Biol. 1995, 253 (3), 493–504.

(71) Ward, S. J.; Gratton, H. E.; Indrayudha, P.; Michavila, C.; Mukhopadhyay, R.; Maurer, S. K.; Caulton, S. G.; Emsley, J.; Dreveny, I. The Structure of the Deubiquitinase USP15 Reveals a Misaligned Catalytic Triad and an Open Ubiquitin-Binding Channel. J. Biol. Chem. 2018, 293 (45), 17362–17374.

(72) Boudreaux, D. A.; Maiti, T. K.; Davies, C. W.; Das, C. Ubiquitin Vinyl Methyl Ester Binding Orients the Misaligned Active Site of the Ubiquitin Hydrolase UCHL1 into Productive Conformation. Proc. Natl. Acad. Sci. U. S. A. 2010, 107 (20), 9117–9122.

(73) Faesen, A. C.; Dirac, A. M. G.; Shanmugham, A.; Ovaa, H.; Perrakis, A.; Sixma, T. K. Mechanism of USP7/HAUSP Activation by Its C-Terminal Ubiquitin-like Domain and Allosteric Regulation by GMP-Synthetase. Mol. Cell 2011, 44 (1), 147–159.

(74) Reily, C.; Stewart, T. J.; Renfrow, M. B.; Novak, J. Glycosylation in Health and Disease. Nat. Rev. Nephrol. 2019, 15 (6), 346–366.

(75) Ramirez, D. H.; Aonbangkhen, C.; Wu, H. Y.; Naftaly, J. A.; Tang, S.; O’Meara, T. R.; Woo, C. M. Engineering a Proximity-Directed O-GlcNAc Transferase for Selective Protein O-GlcNAcylation in Cells. ACS Chem. Biol. 2020, 15 (4), 1059–1066.

(76) Ge, Y.; Ramirez, D. H.; Yang, B.; D’Souza, A. K.; Aonbangkhen, C.; Wong, S.; Woo, C. M. Target Protein Deglycosylation in Living Cells by a Nanobody-Fused Split O-GlcNAcase. Nat. Chem. Biol. 2021, 17 (5), 593–600.

(77) Xiao, H.; Woods, E. C.; Vukojicic, P.; Bertozzi, C. R. Precision Glycocalyx Editing as a Strategy for Cancer Immunotherapy. Proc. Natl. Acad. Sci. U. S. A. 2016, 113 (37), 10304–10309.

(78) Gray, M. A.; Stanczak, M. A.; Mantuano, N. R.; Xiao, H.; Pijnenborg, J. F. A.; Malaker, S. A.; Miller, C. L.; Weidenbacher, P. A.; Tanzo, J. T.; Ahn, G.; Woods, E. C.; Läubli, H.; Bertozzi, C. R. Targeted Glycan Degradation Potentiates the Anticancer Immune Response in Vivo. Nat. Chem. Biol. 2020, 16 (12), 1376–1384.

(79) Ward, C. C.; Kleinman, J. I.; Brittain, S. M.; Lee, P. S.; Chung, C. Y. S.; Kim, K.; Petri, Y.; Thomas, J. R.; Tallarico, J. A.; McKenna, J. M.; Schirle, M.; Nomura, D. K. Covalent Ligand Screening Uncovers a RNF4 E3 Ligase Recruiter for Targeted Protein Degradation Applications. ACS Chem. Biol. 2019, 14 (11), 2430–2440.

(80) Zhang, X.; Crowley, V. M.; Wucherpfennig, T. G.; Dix, M. M.; Cravatt, B. F. Electrophilic PROTACs That Degrade Nuclear Proteins by Engaging DCAF16. Nat. Chem. Biol. 2019, 15 (7), 737–746.

(81) Spradlin, J. N.; Hu, X.; Ward, C. C.; Brittain, S. M.; Jones, M. D.; Ou, L.; To, M.; Proudfoot, A.; Ornelas, E.; Woldegiorgis, M.; Olzmann, J. A.; Bussiere, D. E.; Thomas, J. R.; Tallarico, J. A.; McKenna, J. M.; Schirle, M.; Maimone, T. J.; Nomura, D. K. Harnessing the Anti-Cancer Natural Product Nimbolide for Targeted Protein Degradation. Nat. Chem. Biol. 2019, 15 (7), 747–755.

(82) Luo, M.; Spradlin, J. N.; Boike, L.; Tong, B.; Brittain, S. M.; McKenna, J. M.; Tallarico, J. A.; Schirle, M.; Maimone, T. J.; Nomura, D. K. Chemoproteomics-Enabled Discovery of Covalent RNF114-Based Degraders That Mimic Natural Product Function. Cell Chem. Biol. 2021, 28 (4), 559–566.e15.

(83) Tong, B.; Luo, M.; Xie, Y.; Spradlin, J. N.; Tallarico, J. A.; McKenna, J. M.; Schirle, M.; Maimone, T. J.; Nomura, D. K. Bardoxolone Conjugation Enables Targeted Protein Degradation of BRD4. Sci. Rep. 2020, 10 (1).

(84) He, Y.; Selvaraju, S.; Curtin, M. L.; Jakob, C. G.; Zhu, H.; Comess, K. M.; Shaw, B.; The, J.; Lima-Fernandes, E.; Szewczyk, M. M.; Cheng, D.; Klinge, K. L.; Li, H. Q.; Pliushchev, M.; Algire, M. A.; Maag, D.; Guo, J.; Dietrich, J.; Panchal, S. C.; Petros, A. M.; Sweis, R. F.; Torrent, M.; Bigelow, L. J.; Senisterra, G.; Li, F.; Kennedy, S.; Wu, Q.; Osterling, D. J.; Lindley, D. J.; Gao, W.; Galasinski, S.; Barsyte-Lovejoy, D.; Vedadi, M.; Buchanan, F. G.; Arrowsmith, C. H.; Chiang, G. G.; Sun, C.; Pappano, W. N. The EED Protein–Protein Interaction Inhibitor A-395 Inactivates the PRC2 Complex. Nat. Chem. Biol. 2017 134 2017, 13 (4), 389–395.

(85) Qi, W.; Zhao, K.; Gu, J.; Huang, Y.; Wang, Y.; Zhang, H.; Zhang, M.; Zhang, J.; Yu, Z.; Li, L.; Teng, L.; Chuai, S.; Zhang, C.; Zhao, M.; Chan, H.; Chen, Z.; Fang, D.; Fei, Q.; Feng, L.; Feng, L.; Gao, Y.; Ge, H.; Ge, X.; Li, G.; Lingel, A.; Lin, Y.; Liu, Y.; Luo, F.; Shi, M.; Wang, L.; Wang, Z.; Yu, Y.; Zeng, J.; Zeng, C.; Zhang, L.; Zhang, Q.; Zhou, S.; Oyang, C.; Atadja, P.; Li, E. An Allosteric PRC2 Inhibitor Targeting the H3K27me3 Binding Pocket of EED. Nat. Chem. Biol. 2017 134 2017, 13 (4), 381–388.

(86) Grebien, F.; Vedadi, M.; Getlik, M.; Giambruno, R.; Grover, A.; Avellino, R.; Skucha, A.; Vittori, S.; Kuznetsova, E.; Smil, D.; Barsyte-Lovejoy, D.; Li, F.; Poda, G.; Schapira, M.; Wu, H.; Dong, A.; Senisterra, G.; Stukalov, A.; Huber, K. V. M.; Schönegger, A.; Marcellus, R.; Bilban, M.; Bock, C.; Brown, P. J.; Zuber, J.; Bennett, K. L.; Al-Awar, R.; Delwel, R.; Nerlov, C.; Arrowsmith, C. H.; Superti-Furga, G. Pharmacological Targeting of the Wdr5-MLL Interaction in C/EBPα N-Terminal Leukemia. Nat. Chem. Biol. 2015 118 2015, 11 (8), 571–578.

(87) Varadi, M.; Anyango, S.; Deshpande, M.; Nair, S.; Natassia, C.; Yordanova, G.; Yuan, D.; Stroe, O.; Wood, G.; Laydon, A.; Žídek, A.; Green, T.; Tunyasuvunakool, K.; Petersen, S.; Jumper, J.; Clancy, E.; Green, R.; Vora, A.; Lutfi, M.; Figurnov, M.; Cowie, A.; Hobbs, N.; Kohli, P.; Kleywegt, G.; Birney, E.; Hassabis, D.; Velankar, S. AlphaFold Protein Structure Database: Massively Expanding the Structural Coverage of Protein-Sequence Space with High-Accuracy Models. Nucleic Acids Res. 2021.

(88) Jumper, J.; Evans, R.; Pritzel, A.; Green, T.; Figurnov, M.; Ronneberger, O.; Tunyasuvunakool, K.; Bates, R.; Žídek, A.; Potapenko, A.; Bridgland, A.; Meyer, C.; Kohl, S. A. A.; Ballard, A. J.; Cowie, A.; Romera-Paredes, B.; Nikolov, S.; Jain, R.; Adler, J.; Back, T.; Petersen, S.; Reiman, D.; Clancy, E.; Zielinski, M.; Steinegger, M.; Pacholska, M.; Berghammer, T.; Bodenstein, S.; Silver, D.; Vinyals, O.; Senior, A. W.; Kavukcuoglu, K.; Kohli, P.; Hassabis, D. Highly Accurate Protein Structure Prediction with AlphaFold. Nat. 2021 5967873 2021, 596 (7873), 583–589.

(89) Baek, M.; DiMaio, F.; Anishchenko, I.; Dauparas, J.; Ovchinnikov, S.; Lee, G. R.; Wang, J.; Cong, Q.; Kinch, L. N.; Dustin Schaeffer, R.; Millán, C.; Park, H.; Adams, C.; Glassman, C. R.; DeGiovanni, A.; Pereira, J. H.; Rodrigues, A. V.; Van Dijk, A. A.; Ebrecht, A. C.; Opperman, D. J.; Sagmeister, T.; Buhlheller, C.; Pavkov-Keller, T.; Rathinaswamy, M. K.; Dalwadi, U.; Yip, C. K.; Burke, J. E.; Christopher Garcia, K.; Grishin, N. V.; Adams, P. D.; Read, R. J.; Baker, D. Accurate Prediction of Protein Structures and Interactions Using a Three-Track Neural Network. Science (80-.). 2021, 373 (6557), 871–876.

