## Supplementary information for "Structure-Based Survey of the Human Proteome for Opportunities in Proximity Pharmacology"

**Table S1:** Putative ligandable domains (defined by their Interpro ID)1 and associated list of human protein-modifying enzymes ……………………………………………………..…..…p. S-2

**Table S2**: Ligandable non-catalytic pockets in protein modifying enzymes including acetyltransferases, deacetylases, methyltransferases, demethylases, glycosyltransferases, glycosidases, deubiquitinases, protein kinases and protein phosphatases..……………………………..…………………………………...……p. S-4

**Figure S1**: Recurrent non-catalytic pockets in catalytic domain of Protein arginine methyltransferases...……………………………..…………………………..…...…p. S-5

**References**: ……………………...…………………………………………..……..……p. S-5

**Table S1:** Putative ligandable domains (defined by their Interpro ID)1 and associated list of human protein-modifying enzymes

Acetyltransferases, Deacetylase, Demethylase, Deubiquitinase, Glycosidase, Glycosyltransferase, Methyltransferase, Protein Kinase, Protein Phosphatase

| Domain Name | InterPro ID | Protein names |
|---|---|---|
| Bromo adjacent homology (BAH) domain | IPR001025 | ASH1L |
| Bromo domain | IPR001487, IPR018359 | ASH1L, CREBBP, EP300, KAT2A, KAT2B, KMT2A, TAF1 |
| C1 domain | IPR002219, IPR002420 | ARAF, BRAF, CDC42BPA, CDC42BPB, CDC42BPG, CIT, KSR1, KSR2, PRKCA, PRKCB, PRKCD, PRKCE, PRKCG, PRKCH, PRKCI, PRKCQ, PRKCZ, PRKD1, PRKD2, PRKD3, RAF1, ROCK1, ROCK2, TNS2 |
| C2 domain | IPR002420, IPR000008 | PKN2, PRKCA, PRKCB, PRKCE, PRKCG, PRKCH |
| CARD domain | IPR001315, IPR042147 | RIPK2 |
| EF-hand (associated) domain | IPR013567, IPR002048 | PPEF1, PPEF2, USP32 |
| Immunoglobulin(-like) domain | IPR007110, IPR013783, IPR003599, IPR003598 | ALPK2, ALPK3, AXL, CSF1R, FGFR1, FGFR2, FGFR3, FGFR4, FLT1, FLT3, FLT4, KALRN, KDR, KIT, MERTK, MUSK, MYLK, NTRK1, NTRK2, NTRK3, OBSCN, PDGFRA, PDGFRB, PTPRD, PTPRF, PTPRK, PTPRM, PTPRS, PTPRT, ROR2, SPEG, TEK, TRIO, TTN, TYRO3 |

| | | |
|---|---|---|
| PDZ domain | IPR041489, IPR001478 | CASK, LIMK1, LIMK2, MAST1, MAST2, MAST3, MAST4, PTPN13, PTPN3, PTPN4 |
| POLO box domain | IPR000959, IPR033695 | PLK1, PLK2, PLK3, PLK4 |
| PWWP domain | IPR000313 | NSD1, NSD2, NSD3 |
| Pleckstrin homology domain | IPR001849, IPR041381, IPR043969, IPR033931 | AKT1, AKT2, AKT3, BMX, BTK, CDC42BPA, CDC42BPB, CDC42BPG, CIT, GRK2, GRK3, ITK, JAK1, JAK2, JAK3, KALRN, MAP3K15, MAP3K5, MAP3K6, OBSCN, PDPK1, PHLPP1, PRKD1, PRKD2, PRKD3, ROCK1, ROCK2, TEC, TRIO, TYK2 |
| RCC1-like domain | IPR000408 | NEK8, NEK9 |
| SH2(-like) domain | IPR000980 | ABL1, ABL2, BLK, BMX, BTK, CSK, FER, FES, FGR, FRK, FYN, HCK, ITK, JAK1, JAK2, JAK3, LCK, LYN, MATK, PTK6, PTPN11, PTPN6, SOCS1, SOCS2, SOCS3, SOCS4, SOCS5, SOCS6, SOCS7, SRC, SRMS, SYK, TEC, TNS2, TXK, TYK2, YES1, ZAP70 |
| SH3 domain | IPR001452, IPR035750 | ABL1, ABL2, BLK, BTK, CASK, CSK, FGR, FRK, FUT8, FYN, HCK, ITK, KALRN, LCK, LYN, MAP3K10, MAP3K11, MAP3K21, MAP3K9, MATK, OBSCN, PRMT2, PTK6, SRC, SRMS, TEC, TNK1, TNK2, TRIO, TXK, UBASH3B, YES1 |

| | | |
|---|---|---|
| SWIRM domain | IPR007526 | KDM1A, KDM1B, MYSM1 |
| TIM barrel domain | IPR035247 | PRMT5 |
| Tetratricopeptide repeat | IPR001440, IPR019734, IPR013026, IPR006597 | EEF2K, KDM6A, OGT, PPP5C, PRMT9, TMTC1, TMTC2, TMTC3, TMTC4, UTY |
| Tudor domain | IPR041292, IPR002999 | KDM4A, KDM4B, KDM4C, SETDB1, STK31 |
| Ubiquitin carboxyl-terminal hydrolase, C-terminal | IPR029346 | USP7, USP11, USP15 |
| WD 40 repeat | IPR001680, IPR018391, IPR018391, IPR019775, IPR017986, IPR036322, IPR015943 | GTF3C4, LRRK1, LRRK2, MET, MST1R, PIK3R4 |
| WW domain | IPR001202 | GALNT9, SETD2 |
| Zinc finger, UBP-type | IPR001607 | HDAC6, USP3, USP5, USP13, USP16, USP20, USP22, USP33, USP44, USP45, USP49, USP51 |

**Table S2: Ligandable non-catalytic pockets in protein modifying enzymes including acetyltransferases, deacetylases, methyltransferases, demethylases, glycosyltransferases, glycosidases, deubiquitinases, protein kinases and protein phosphatases.**

Each row represents a non-catalytic ligandable pocket and the pocket is labeled according to protein family and domain location in the column ‘Label’. Confidence level indicates whether a potent ligand is bound to the pocket in the specified protein (confidence level 1), a peptide or >10nM ligand bound to the pocket in the specified protein (confidence level 2), a pocket in a domain demonstrated to be ligandable in other proteins (confidence level 3) or pocket that meets the ligandability criteria but has no reported ligands in the protein of interest or homologues (confidence level 4). Pockets with similar label and color (column ‘Color group’) represent a recurrent pocket location in a domain within the protein family. ‘Pocket details’ column contains additional information about the pocket, for example pocket names. Pockets are labelled with a PDB code, gene name and Uniprot ID. For each pocket, properties such as pocket residues, reactive cysteine residues, InterPro code and InterPro domain name are labeled according to the chain in the PDB structure in which the pocket is identified (for example, pocket residues a/123,124 are residues 123 and 124 in chain a). Pocket residues are the residues lining the pocket. (Residues within 2.8Å distance from the pocket mesh generate by ICM). Reactive cysteine residues are cysteine residues lining the pocket and predicted reactivity of cysteine residue is calculated using the ReactiveCys module of ICM. (The reactivity score is based on a weighted function of the energy and distance data reported in the results table). The columns ‘InterPro code’ and ‘InterPro domain name’ provide the InterPro code and corresponding InterPro domain name of the domain where the pocket is located. Distance to catalytic residues is calculated in ICM (Å). N/A denotes absence of catalytic residues in the structure. Pocket properties including, ‘Volume’, ‘Area’, ‘Hydrophobicity’, ‘Buriedness’ and ‘DLID’ are calculated by the icmPocketfinder module of ICM.

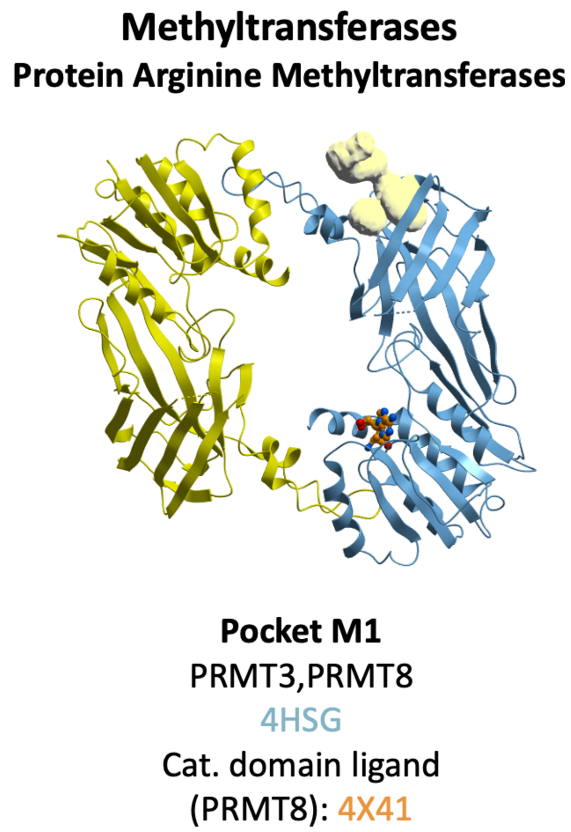


**Figure S1. Recurrent non-catalytic pockets in catalytic domain of Protein arginine methyltransferases.** Protein Arginine methyltransferase (blue) bound to catalytic inhibitor (orange). Reference structure: PDB 4HSG[2]; reference catalytic inhibitor (orange): PDB 4X41[3].

References

(1) Blum, M.; Chang, H. Y.; Chuguransky, S.; Grego, T.; Kandasaamy, S.; Mitchell, A.; Nuka, G.; Paysan-Lafosse, T.; Qureshi, M.; Raj, S.; Richardson, L.; Salazar, G. A.; Williams, L.; Bork, P.; Bridge, A.; Gough, J.; Haft, D. H.; Letunic, I.; Marchler-Bauer, A.; Mi, H.; Natale, D. A.; Necci, M.; Orengo, C. A.; Pandurangan, A. P.; Rivoire, C.; Sigrist, C. J. A.; Sillitoe, I.; Thanki,

N.; Thomas, P. D.; Tosatto, S. C. E.; Wu, C. H.; Bateman, A.; Finn, R. D. The InterPro Protein Families and Domains Database: 20 Years On. *Nucleic Acids Res.* 2021, *49* (D1), D344–D354.

(2) Liu, F.; Li, F.; Ma, A.; Dobrovetsky, E.; Dong, A.; Gao, C.; Korboukh, I.; Liu, J.; Smil, D.; Brown, P. J.; Frye, S. V.; Arrowsmith, C. H.; Schapira, M.; Vedadi, M.; Jin, J. Exploiting an Allosteric Binding Site of PRMT3 Yields Potent and Selective Inhibitors. *J. Med. Chem.* 2013, *56* (5), 2110–2124.

(3) Lee, W. C.; Lin, W. L.; Matsui, T.; Chen, E. S. W.; Wei, T. Y. W.; Lin, W. H.; Hu, H.; Zheng, Y. G.; Tsai, M. D.; Ho, M. C. Protein Arginine Methyltransferase 8: Tetrameric Structure and Protein Substrate Specificity. *Biochemistry* 2015, *54* (51), 7514–7523.
